# Leveraging High-Throughput Proteomics and AI-Based Protein Folding to Accelerate VAV1 Molecular Glue Discovery

**DOI:** 10.1101/2025.06.08.658535

**Authors:** Hanfeng Lin, Xin Yu, Haiyang Zheng, Ran Cheng, Min Zhang, Xiaoli Qi, Yen-Yu Yang, Shengmin Zhou, Rui Qi, Ly Le, Andrea Bortolato, Semen Yesylevskyy, Alan Nafiiev, Xing Che, Jin Wang

**Affiliations:** The Verna and Marrs McLean Department of Biochemistry and Molecular Pharmacology, Baylor College of Medicine, Houston, Texas 77030, United States; Center for NextGen Therapeutics, Baylor College of Medicine, Houston, Texas 77030, United States.; Graduate School of Biomedical Sciences, Baylor College of Medicine, Houston, Texas 77030, United States.; Thermo Fisher Scientific, San Jose, California 95134, United States.; YDS Pharmatech, Inc., Albany, New York 12226, United States.; SandboxAQ, Palo Alto, California 94301, United States.; Receptor.AI Inc., 223 Concord Turnpike, Cambridge, Massachusetts 02140, United States.; Department of Molecular and Cellular Biology, Baylor College of Medicine, Houston, Texas 77030, United States.

## Abstract

This study reports a series of CRBN molecular glues that selectively induce proteasomal degradation of VAV1, a hematopoietic-specific signaling protein and key target in hematological malignancies and autoimmune diseases. Using unbiased global proteomics, we identify phenyl-glutarimide derivatives (e.g., NGT-201-12) as effective VAV1 degraders that engage its C-terminal SH3 domain (SH3-2). We elucidate a non-canonical RT-loop degron (RDxS motif, residues 796–799) in SH3-2, distinct from previously characterized G-loop degrons, supported by the physics- and AI-driven GluePlex workflow and site-directed mutagenesis. Applying Free Energy Perturbation (FEP) to predicted ternary structures yields cooperativity metrics that correlate with degradation potency, overcoming limitations of standard docking and enabling prospective ranking of analogs—even from weak initial binders. We further show that conformational restriction via halogen substitution (e.g., NGT-201-18) potentiates degradation, a principle rationalized by DFT calculations, while comprehensive SAR studies chart a roadmap for next-generation degraders. Dose-response proteomics confirms VAV1 as the primary target and reveals LIMD1, bearing a canonical G-loop, as an off-target, showing that one glue can engage disparate degrons. These findings expand the landscape of CRBN neosubstrate recognition and offer a therapeutic strategy for targeting VAV1.

## Introduction

The ubiquitin-proteasome system (UPS) is a fundamental cellular machinery responsible for maintaining protein homeostasis through the selective degradation of a vast array of intracellular proteins, including those that are misfolded, damaged, or naturally short-lived ^1^. This intricate process is critical for normal cellular function, and its dysregulation is implicated in numerous human diseases^1^. In recent years, the therapeutic strategy of Targeted Protein Degradation (TPD) has gained prominence, aiming to eliminate disease-causing proteins by hijacking the UPS ^2–7^. Among TPD approaches, molecular glue degraders have emerged as a compelling class of small molecules ^8^. These agents function by inducing or stabilizing the protein-protein interaction (PPI) between an E3 ubiquitin ligase and a specific target protein (a neosubstrate), which is then polyubiquitinated and subsequently degraded by the proteasome ^9–13^.

Cereblon (CRBN), a substrate receptor for the Cullin-RING E3 ubiquitin ligase 4 (CRL4^CRBN^) complex, is a well-established E3 ligase that can be effectively modulated by molecular glues ^8,10,14^. The therapeutic utility of CRBN modulation was first highlighted by immunomodulatory drugs (IMiDs)—thalidomide, lenalidomide, and pomalidomide ^9,15,16^. These drugs bind to CRBN, altering its substrate specificity to promote the degradation of neosubstrates such as the lymphoid transcription factors IKZF1 and IKZF3. These canonical substrates typically engage CRBN via a structural motif known as the G-loop (a β-hairpin turn containing a conserved glycine residue), which is crucial for their anti-myeloma activity ^9,11^. The profound clinical success of IMiDs has not only validated CRBN as a druggable E3 ligase but has also catalyzed extensive research efforts aimed at discovering novel CRBN modulators capable of degrading a wider array of pathogenic proteins ^17–19^.

The discovery of novel molecular glues, however, remains a significant challenge due to the indirect nature of their mechanism, which complicates rational design efforts ^8,13^. Historically, many glues were found serendipitously ^20^, through target-driven screening using reporter assay system ^21^, or comparative phenotypic screening in E3-deficient cell models ^22^. Modern approaches increasingly rely on systematic strategies, especially advanced proteomic techniques for unbiased neosubstrate identification and selectivity profiling. Degradation-based global proteomics enables direct identification of degradable neosubstrates ^12^. Proximity-based proteomics helps identify transient protein-protein interaction mediated by lead compounds ^23,24^. Enrichment-based chemoproteomics, for instance, allows for the mapping of proteins recruited to CRBN by specific compounds ^18^. Computational tools like YDS-GlueFold is combining diffusion model with statistic mechanics for predicting ternary complex structures and guiding molecular design ^25^, while FEP+ binding free energy calculation in the ternary system for explaining cooperativity and target selectivity ^26^.

VAV1, a hematopoietic-specific signaling protein, is a key regulator of immune cell development and function ^27^. It acts as both a guanine nucleotide exchange factor (GEF) for Rho/Rac family GTPases and as an adaptor protein, playing critical roles in T-cell receptor (TCR) and B-cell receptor (BCR) signaling pathways that govern calcium mobilization, ERK activation, cytoskeletal rearrangements, and gene transcription ^28^. Dysregulation of VAV1, through mutations or aberrant expression, has been linked to various hematological malignancies including T-cell lymphomas, and autoimmune diseases ^29^. Given its multifaceted role in disease and its hematopoietic-specific expression, VAV1 presents an attractive therapeutic target ^30^. Traditional inhibitors might only partially modulate VAV1’s functions, whereas its complete elimination via targeted protein degradation could offer a more comprehensive therapeutic effect by ablating both its catalytic and scaffolding activities ^28^.

In this study, we report the identification and characterization of a series of CRBN molecular glues that selectively target VAV1 for degradation. Utilizing an unbiased, deep global proteomics-based approach enabled by Thermo Orbitrap Astral Mass Spectrometer, we identified NGT-201-11, a phenyl-glutarimide derivative, and NGT-201-12, an indole-modified analog with enhanced potency, as effective VAV1 degraders. Mechanistic investigations employing a luminescence-based HiBiT-RR protein quantification system ^31^ with VAV1 truncation mutants localized the essential degron to the C-terminal SH3 domain (SH3-2) of VAV1. We developed the physics- and AI-driven GluePlex pipeline and computationally modeled the CRBN:NGT-201-12:VAV1-SH3-2 ternary complex, predicting critical interactions involving VAV1 R796 and the RT loop of the SH3-2 domain. While established structural biology techniques like CryoEM and X-ray crystallography are considered the gold standards for determining molecular structures, they typically require highly optimized compounds and conditions, especially for resolving complex assemblies ^11^. In contrast, our GluePlex provides a significant advantage by enabling the prediction of ternary complex structures during the early stages of drug discovery when such structural information can be most impactful for guiding compound design and optimization. Site-directed mutagenesis studies functionally validated this predicted RT loop (a variable surface loop connecting β-strands within the SH3 domain fold), specifically an RDxS motif (residues 796-799), as a non-canonical degron essential for NGT-201-12-mediated degradation. This finding is significant as it deviates from the canonical G-loop degrons typically recognized by CRBN ^8^. Further chemical optimization, including conformational restriction via halogen substitution (e.g., NGT-201-17, NGT-201-18), led to potentiated VAV1 degradation. Dose-response proteomics-based structure-activity relationship analyses not only confirmed VAV1 as the primary target but also unexpectedly identified a G-loop-degron-possessing LIM Domain Containing 1 (LIMD1) as an off-target for some compounds in this series, This highlights the capacity of a single molecular glue to engage distinct degron motifs on canonical G-loop and noncanonical G-loop independent neosubstrates, underscoring the critical role of comprehensive proteomic profiling in molecular glue discovery and optimization.

## Results

### Proteomics Screening Identifies VAV1 as a Neosubstrate of CRBN Glue

To investigate avenues of CRBN-mediated protein degradation, an unbiased, quantitative mass spectrometry-based proteomics approach was utilized to screen a phenyl-glutarimide centric library ^32,33^, analyzing cellular proteomes after treatment with candidate CRBN-based molecular glue compounds (**Figure 1A, B**). From this library, NGT-201-11 and NGT-201-12, possessing similar substitution patterns on the core phenyl-glutarimide scaffold, were concurrently identified as compounds capable of inducing VAV1 degradation, albeit to different degrees. Cellular treatment with NGT-201-11, which features a phenyl group modification, led to a modest yet statistically significant decrease in VAV1 protein levels compared to the vehicle controls (**Figure 1C**). NGT-201-12, an analog where the phenyl moiety is replaced by an indole group, demonstrated a more pronounced degradation of VAV1 (**Figure 1D**). The degradation of VAV1 by NGT-201-12 was subsequently confirmed through Western blot analysis, while the potency of NGT-201-11 was too weak to distinguish on the blot (**Supplementary Figure 1A-B**). The degradation of VAV1 by NGT-201-12 can be rescued by blocking the E1 (TAK-243), NEDDylation (MLN-4924), proteasome (carfilzomib), and CRBN (pomalidomide) activities or through CRBN-knockout (**Supplementary Figure 1E-F**). Degradation kinetic experiment showed that dose-dependent degradation could be observed as short as 2 h of NGT-201-12 treatment (**Supplementary Figure 1C**). The VAV1 mRNA level was not affected by NGT-201-12 treatment (**Supplementary Figure 1H**). These findings validated the potential of this chemotype and highlighted NGT-201-12 as a targeted-protein degradation candidate for more detailed mechanistic studies and further optimization.

**Figure 1.**
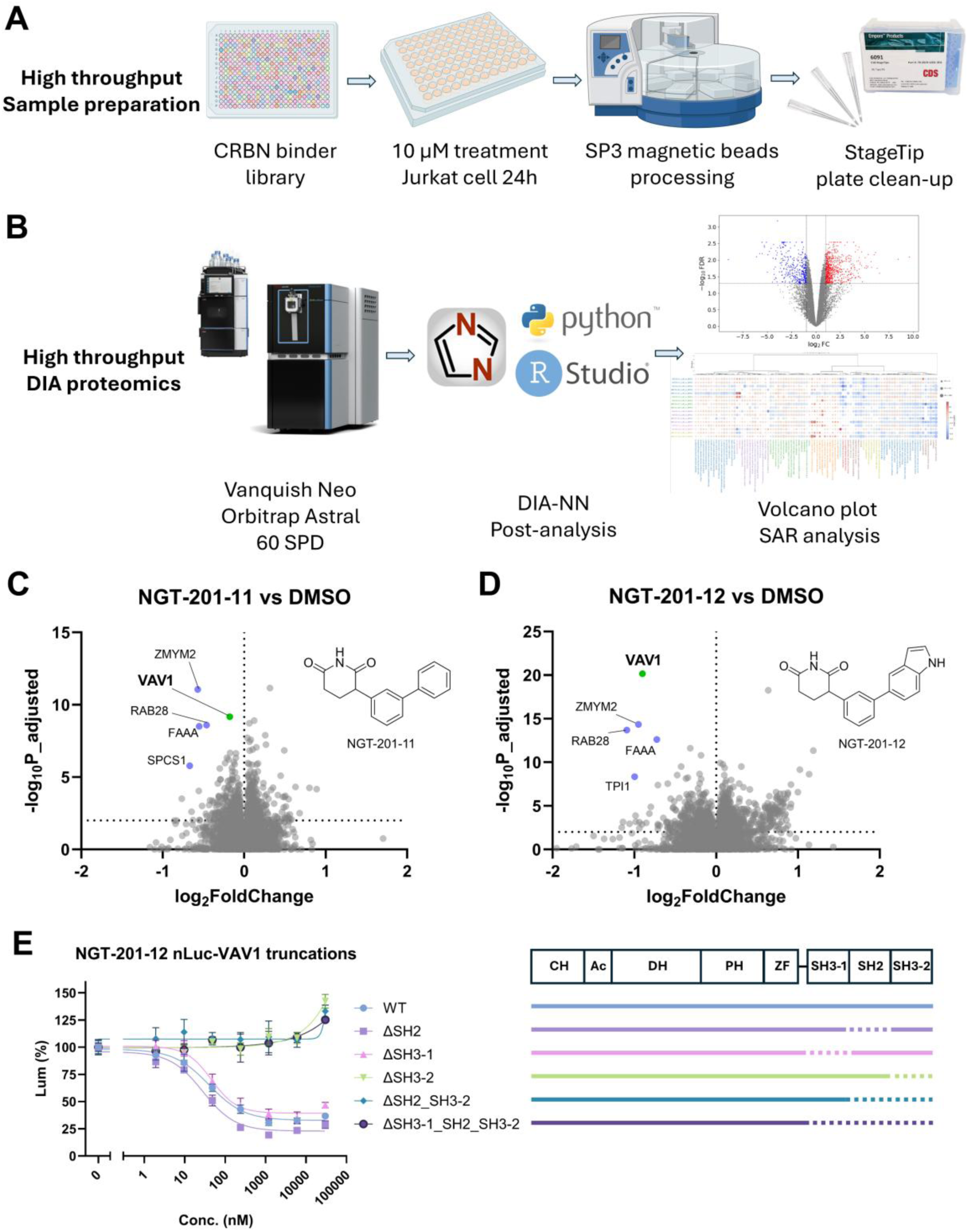
Proteomics Screening Identifies VAV1 as a Neosubstrate of CRBN Molecular Glues and Localization of the Degron to the C-terminal SH3 Domain. **(A)** Schematic representation of the high-throughput DIA proteomics workflow employed to identify potential CRBN neosubstrates. Created in BioRender. Lin, H. (2025) https://BioRender.com/7rhcf1h. (**B**) Overview of the DIA-NN data processing and statistical analysis pipeline. SPD, sample-per-day. Created in BioRender. Lin, H. (2025) https://BioRender.com/7rhcf1h. The R logo is © 2016 The R Foundation. The logo is licensed under the terms of the Creative Commons Attribution-ShareAlike 4.0 International license (CC-BY-SA 4.0 https://creativecommons.org/licenses/by-sa/4.0/). (**C, D**) Volcano plots depicting protein abundance changes in Jurkat cells treated with 10 μM NGT-201-11 (**C**) or NGT-201-12 (D) for 24 hours compared to DMSO control. VAV1 is green highlighted. Other significantly downregulated proteins are in blue. Differentially abundant proteins were identified using a moderated t-test (limma) with two-sided p-values; p-values were adjusted for multiple comparisons using the Benjamini–Hochberg procedure to control the false discovery rate (FDR). (**E**) HiBiT-based dose-response curves illustrating the degradation of full-length VAV1 (WT) and various truncation mutants (ΔSH2, ΔSH3-1, ΔSH3-2, ΔSH2_SH3-2, ΔSH3-1_SH2_SH3-2) upon 24 hours treatment with increasing concentrations of NGT-201-12 in Hela cells. Loss of the C-terminal SH3 domain (ΔSH3-2) abolishes NGT-201-12-induced degradation. Data are presented as mean ± SD from three independent experiments.

VAV1 exhibits a modular structure comprising tandem SH3-1_SH2_SH3-2 domains on its C-terminus integral to its signaling capacity. To precisely delineate the VAV1 region mediating interaction with the CRBN:NGT-201-12 complex, a luminescence-based protein quantification approach utilizing the HiBiT-RR system was employed to avoid tag-mediated degradation artifacts ^31^. We fused HiBiT-RR to N- or C-terminus of full length VAV1 and observed no difference in degradation potency of NGT-201-12 (**Supplementary Figure 1G**). A series of VAV1 truncation mutants fused with C-terminus HiBiT-RR peptide tag was then generated. Consistent with expectations, cells expressing full-length VAV1-HiBiT-RR constructs demonstrated robust, dose-dependent reduction in luminescence signal following NGT-201-12 treatment. When truncating domains from the C-terminus, we discovered that removal of the SH3 domain on the C-terminus (SH3-2) is sufficient and necessary to abrogate NGT-201-12-induced VAV1 degradation (**Figure 1E, Supplementary Figure 1I**). These findings localize the essential degron recognized by the CRBN:NGT-201-12 complex to the C-terminal SH3-2 domain of VAV1.

### Ternary Complex Modeling Predicts Key Molecular Interactions

Molecular glues function by inducing de novo protein-protein interactions (PPIs) between an E3 ligase and a target protein. Although ligand-dependent, these assemblies frequently anchor upon intrinsic surface geometries and energetic hotspots characteristic of natural PPI interfaces ^14,34,35^. To capitalize on this principle, we developed GluePlex, an integrated structural modeling pipeline designed to predict the ternary complex of CRBN:NGT-201-12:VAV1 SH3-2. GluePlex takes advantage of open-source geometric deep learning, physics-based docking, and templated co-folding to maximize reproducibility (**Figure 2A**). First, we utilized the open-source geometric deep learning model PeSTo ^36^ to predict protein-protein interaction hotspots on the unbound surfaces of CRBN and VAV1 SH3-2 domains. These predicted contact zones served as biological restraints for HADDOCK docking ^37^, generating a diverse ensemble of binary protein complex templates. Subsequently, using one representative conformation from each HADDOCK cluster as structural scaffolds, we employed the open-source Boltz-2 ^38^ to perform templated co-folding with NGT-201-12, enabling the precise, induced-fit positioning of the small molecule within the interface. UMAP analysis of the resulting 120 co-folded structures revealed distinct conformational populations even starting from the same clustered templates (**Figure 2B**). We prioritized the cluster exhibiting the highest interface confidence metrics, which corresponded to a stable complex assembly compatible with the ligand geometry. Consequently, the top scored structure from this cluster (ipTM = 0.94, chainpair-ipTM = 0.89) was selected as the functional ternary complex model (**Figure 2C**).

**Figure 2.**
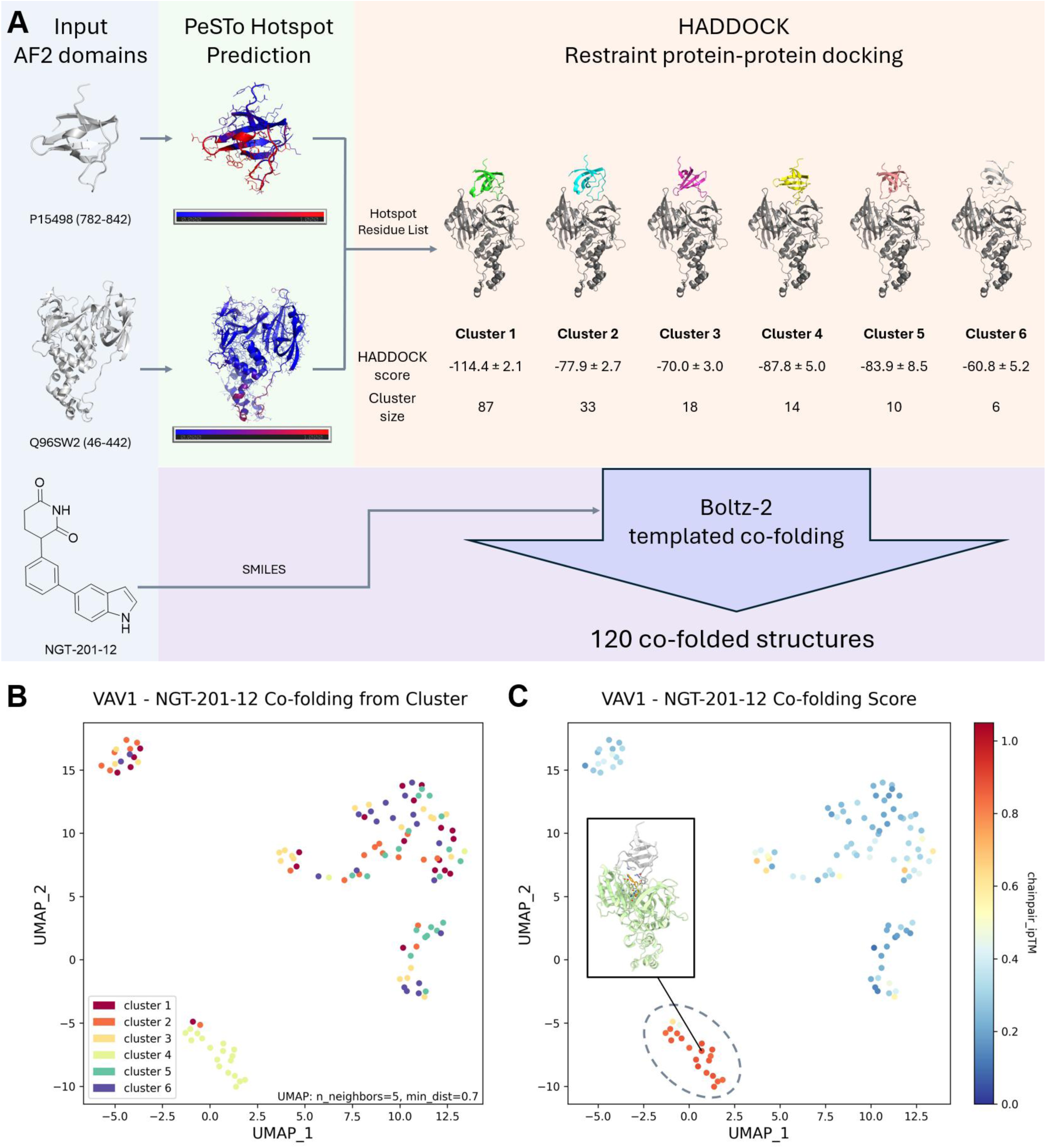
GluePlex workflow for structural prediction of the CRBN:NGT-201-12:VAV1 ternary complex. **(A)** Schematic of the integrated open-source modeling pipeline. AlphaFold2 structures of the VAV1 SH3-2 domain (P15498) and CRBN (Q96SW2) were analyzed by the geometric deep learning model PeSTo to predict protein-protein interaction hotspots (red for hotspots). These predicted active residues served as restraints for HADDOCK docking, yielding diverse binary complex clusters (Clusters 1–6) ranked by HADDOCK score. Representative structures from these clusters were used as templates for Boltz-2 co-folding with the NGT-201-12 ligand to generate 120 ternary models. **(B)** UMAP visualization of the conformational landscape of the 120 co-folded ternary structures. Each point represents a generated model, colored by the HADDOCK cluster of origin, showing distinct structural populations. **(C)** Model selection and quality assessment. The UMAP embedding is colored by the chainpair_ipTM confidence score between VAV1 chain and CRBN chain. The high-confidence models (red) form a distinct cluster (originating largely from HADDOCK Cluster 4), identifying the predicted bioactive conformation (ipTM = 0.94) used for experimental validation.

The resultant structural model furnished valuable atomic-level hypotheses concerning the binding interface (**Figure 3A, Supplementary Data 1**). The model indicated that the glutarimide part of NGT-201-12 occupies the canonical IMiD binding site on CRBN, orienting its indole moiety outwards (**Figure 3B, D**). This exposed indole group on NGT-201-12 is predicted to form a composite interface engaging a specific region on the VAV1 SH3-2 domain surface, exemplified by the cation-π interaction between VAV1 R796 and the indole ring in NGT-201-12 (**Figure 3B, D**). Furthermore, it highlighted potentially crucial direct protein-protein interactions (PPIs) between CRBN and VAV1 SH3-2 domain RT loop (residues 796-799, connecting β-strands C and D of the SH3 fold) that could contribute to the stabilization of the ternary complex assembly (**Figure 3C**).

**Figure 3.**
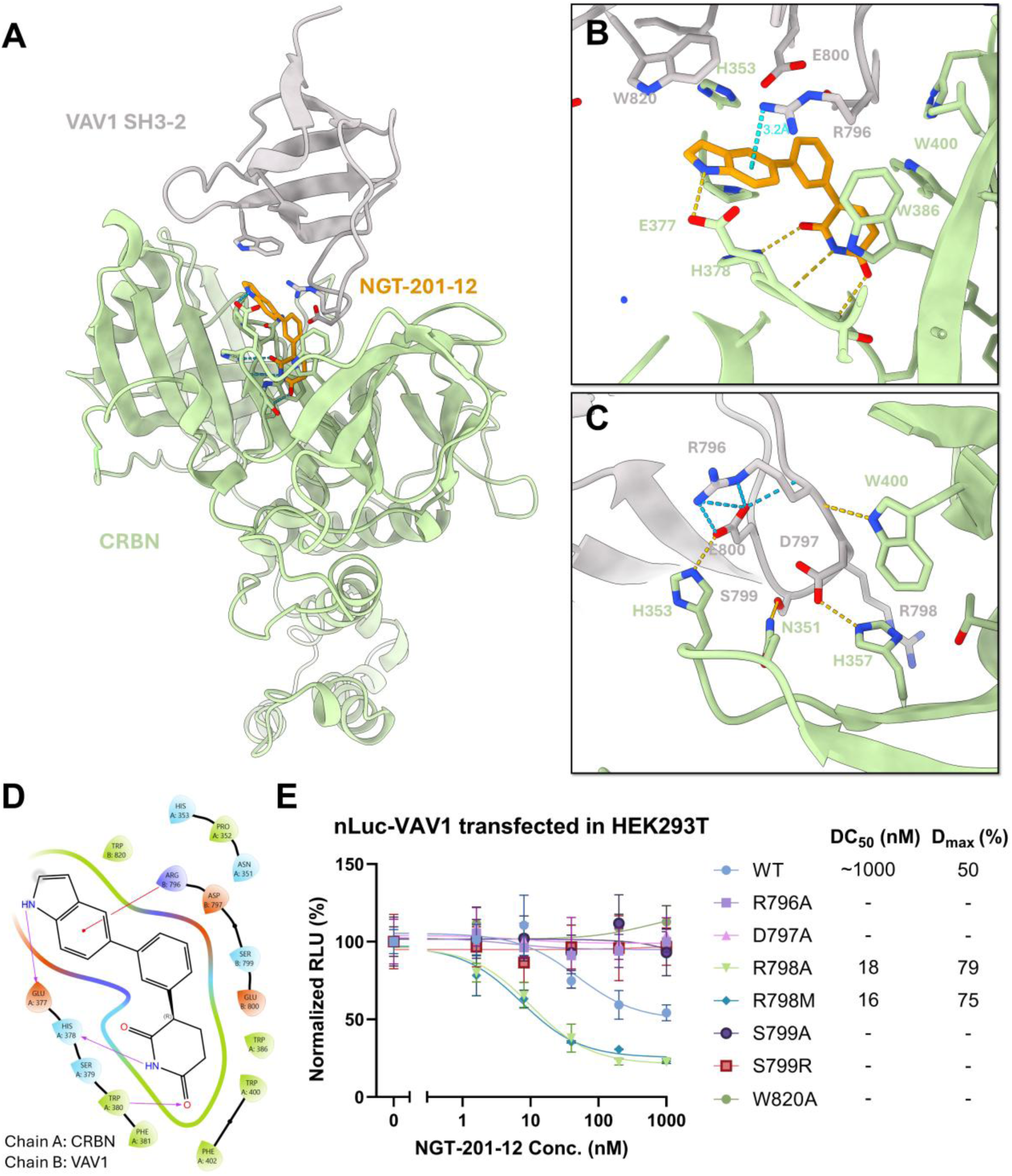
Ternary Complex Modeling and Mutagenesis Identify a Non-Canonical RT Loop Degron in VAV1 SH3-2. **(A)** Computational model of the NGT-201-12-induced ternary complex formed between CRBN and the VAV1 SH3-2 domain. NGT-201-12 is shown in orange, CRBN in light green, and VAV1 SH3-2 in gray. **(B)** Close-up view of the predicted binding interface among VAV1 SH3-2, NGT-201-12 and CRBN, highlighting potential cation-π interaction (cyan dashed lines) and hydrogen bonds (yellow dashed lines). Key CRBN residues involved in the interaction are labeled. **(C)** Close-up view of the predicted interaction interface between NGT-201-12 and the VAV1 SH3-2 domain RT loop region, illustrating potential hydrogen bonds (cyan dashed lines) and hydrophobic interactions (yellow dashed lines). Key VAV1 SH3-2 residues within the RT loop are labeled. **(D)** Ligand interaction diagram generated from the computational model, depicting the predicted interactions between NGT-201-12 and residues in CRBN and VAV1 SH3-2. **(E)** Dose-response curves showing the effect of NGT-201-12 on the degradation of wild-type (WT) nLuc-VAV1 and various point mutants within the predicted CRBN/NGT-201-12 interaction interface in HEK293T cells. Data are presented as mean ± SD from two independent experiments. The table insert summarizes the DC_50_ and D_max_ values for each mutant.

To functionally validate the interface predicted by the GluePlex model, the putative PPI surface on the VAV1 SH3-2 domain was systematically interrogated via site-directed mutagenesis. Residues situated within surface-exposed loops, particularly those predicted to mediate contact with CRBN or NGT-201-12, were mutated, primarily to alanine. The impact of these mutations on compound-induced degradation was subsequently quantified using the nano-luciferase (nLuc)-VAV1 reporter system. Evaluation of mutants treated with NGT-201-12 revealed that mutations within the RT-loop exerted the most destructive effects on degradation efficiency (**Figure 3E, Supplementary Figure 1J**). Specifically, alanine mutation of the core RDxS motif residues—Arg796 (R796A), Asp797 (D797A), Ser799 (S799A) or S799R (mimicking the corresponding residue in VAV2/3)—resulted in a loss of degradation potency, effectively abrogating the dose-response relationship observed with wild-type VAV1 (**Figure 3E**). Conversely, mutation of Arg798 to alanine (R798A) or methionine (R798M, mimicking the corresponding residue in VAV2/3), paradoxically enhanced degradation potency and D_max_ (**Figure 3E**). The enhanced degradation observed with the R798A mutant is rationalized by molecular dynamics (MD) simulations, which identify the wild-type R798 as a steric impediment to ternary complex formation. The simulations show that the large, flexible guanidinium side chain of R798 samples multiple conformations that clash with the CRBN surface (**Supplementary Figure 2D, F, G**). In contrast, the smaller alanine side chain in the R798A mutant completely avoids this steric hindrance (**Supplementary Figure 2E, F, G**). Although wild-type R798 can form transient hydrogen bonds with CRBN residues Y355 and S420, these are infrequent and insufficient to overcome the destabilizing steric effect during the ternary complex formation process (**Supplementary Figure 2A, B**). Consequently, the R798A mutation removes this key impediment, allows for a more stable RT-loop conformation indicated by lower RMSF values of residue 796 and 798 in R798A mutant than WT (**Supplementary Figure 2C**), therefore promotes a more productive protein-protein interface for subsequent degradation. All these computational results are consistent with the experimental result that the R798A mutation leads to stronger VAV1:molecular glue:CRBN ternary complex formation (**Figure 4D**), which further supports our hypothesis that R798A mutant is more prone to being degraded due to easier ternary complex formation. Mutations at CRBN:VAV1 interface outside the immediate RT loop, such as Q818A, exhibited more moderate effects on NGT-201-12-induced degradation (**Supplementary Figure 1K**), whereas W820A mutation, predicted to be at the VAV1:NGT-201-12 interface, significantly reduced degradation potency (**Figure 3E**).

**Figure 4.**
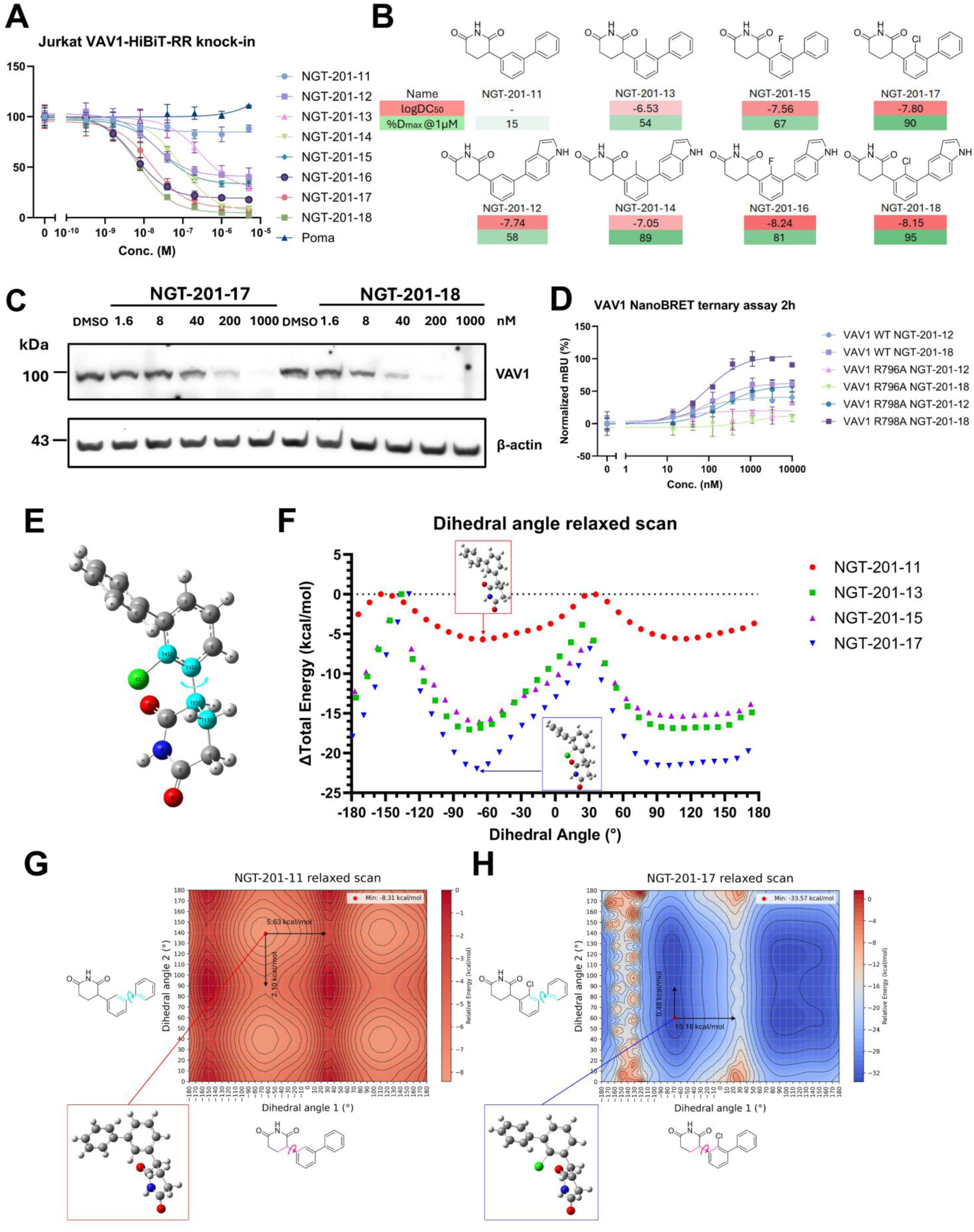
Conformational Restriction via Halogen Substitution Potentiates Degrader Activity. **(A)** Dose-response curves for the degradation of VAV1-HiBiT-RR in Jurkat cells treated with various NGT-201 compounds or pomalidomide for 24 hours. Data are presented as mean ± SD from three independent experiments. **(B)** Chemical structures of the tested NGT-201 compounds, with corresponding logDC_50_ and D_max_ values (at 1 μM) indicated by a color gradient (green for higher D_max_, red for stronger potency or lower logDC_50_). **(C)** Western blot analysis confirming the dose-dependent degradation of endogenous VAV1 by 24 hours NGT-201-17 and NGT-201-18 treatment in Jurkat cells. β-actin serves as a loading control. The blot is representative of three independent experiments. **(D)** Lysate-based NanoBRET VAV1:degrader:CRBN ternary complex assay with nLuc-VAV1 full length WT, R796A, or R798A mutant. Data are presented as mean ± SD from three independent experiments. **(E)** Representative optimized 3D structure of NGT-201-17 obtained from DFT calculations, highlighting the dihedral scanning atoms. **(F)** Dihedral angle relaxed scan showing the calculated potential energy profile for rotation around the bond connecting the phenyl ring and the glutarimide ring for NGT-201-11 (red), −13 (green), −15 (purple), and −17 (blue). NGT-201-17 exhibits a higher rotational energy barrier, suggesting reduced conformational flexibility. The insets depict representative conformations at different dihedral angles. **(G, H)** 2D potential energy surface relaxed scanning both rotatable bonds on NGT-201-11 (G) and −17 (H)

Collectively, these functional mutagenesis data strongly implicate the RT loop, and specifically the RDxS sequence (residues 796-799), as the critical, non-canonical degron within the VAV1 SH3-2 domain essential for mediating the interaction with the CRBN-molecular glue complex. This finding stands in contrast to the canonical G-loop degrons identified in previously characterized IMiD neosubstrates and underscores the capacity of CRBN-glue complexes to recognize diverse structural motifs for neosubstrate recruitment.

### Conformational Restriction via Halogen Substitution Potentiates Degrader Activity

According to the ternary complex model, NGT-201-12 takes a specific torsion angle to extend the indole ring to R798 for maximizing the cation-π interaction. It was hypothesized that the inherent conformational flexibility of NGT-201-12, specifically the rotation around the single bond linking the glutarimide-bearing phenyl ring and the indole moiety, might constitute a potential liability by increasing the entropic cost associated with ternary complex formation ^39,40^. To investigate whether molecular rigidification could enhance potency through increasing the torsional barrier ^39^, small substituents were introduced onto the phenyl ring proximal to the glutarimide moiety to sterically impede free rotation. Methyl (NGT-201-13/14), fluoride (NGT-201-15/16), and chloride (NGT-201-17/18) substitutions were evaluated (**Figure 4A, B**). Assessment of these analogs demonstrated a significant potentiation of VAV1 degradation activity for the chlorinated derivatives, particularly significantly improved maximum degree of VAV1 degradation (D_max_). Both NGT-201-17 (phenyl analog) and NGT-201-18 (indole analog) displayed improved VAV1 degradation relative to their respective non-halogenated precursors, NGT-201-11 and NGT-201-12 (**Figure 4A**), but with no better CRBN binding measured by CRBN NanoBRET target engagement assay (**Supplementary Figure 3**). The VAV1 degradation induced by NGT-201-17 and −18 are further confirmed by immunoblotting against the endogenous VAV1 in Jurkat cells (**Figure 4C**). The degradation mechanism and kinetics of NGT-201-18 was also confirmed to be similar as NGT-201-12 (**Supplementary Figure 1B-F**). The critical dependence on the RDxS motif was also confirmed in NGT-201-17 and NGT-201-18, where mutations R796A, D797A, and S799A consistently led to substantial resistance to degradation (**Supplementary Figure 4**). NGT-201-18 also induces stronger ternary complex formation than NGT-201-12 in the NanoBRET ternary complex experiment, while R796A mutants showed no ternary complex formation for both compounds (**Figure 4D**).

To establish a more rigorous, physics-based rationale for the observed potentiation, Density Functional Theory (DFT) calculations were conducted to compute the potential energy profile associated with rotation around the critical dihedral angle connecting the phenyl and glutarimide rings. We evaluated the highly flexible unsubstituted core (NGT-201-11) alongside its methyl-(NGT-201-13), fluoro-(NGT-201-15), and chloro-substituted (NGT-201-17) counterparts (**Figure 4E, F**). These calculations revealed a clear gradient of conformational restriction. The chlorinated NGT-201-17 exhibits the deepest energetic well and highest rotational barrier, while the fluoro and methyl substitutions impose intermediate torsional barriers relative to the unrestricted NGT-201-11 (**Figure 4F**). Although fluoro and methyl differ in van der Waals radius (∼1.47 vs ∼2.0 Å), the methyl group can rotate its three hydrogens to minimize steric clashes with the adjacent glutarimide carbonyl and therefore leaks conformational energy. The rotatable bond between two phenyl rings, however, is largely permissive for free rotation (**Figure 4G-H**). Molecular dynamics simulations of NGT-201-11/12/17/18 in complex with CRBN also show that the chlorinated compounds have narrower windows of dihedral angle than the non-chlorinated counterparts (**Supplementary Figure 5**). Interestingly, while the methyl analog (NGT-201-13) exhibits a similar torsional energy landscape to the fluoro analog (NGT-201-15), its cellular degradation potency is inferior. This discrepancy is explained by evaluating primary E3 ligase engagement. CRBN NanoBRET binary assay (**Supplementary Figure 3**) demonstrated that the bulkier methyl substitution severely abrogates binary binding to CRBN (IC50 = 883 nM) compared to the fluoro analog (IC50 = 101 nM). Thus, while conformational restriction minimizes the entropic penalty of ternary assembly, successful degradation requires a delicate balance that does not fatally compromise the initial binary engagement with the CRBN pocket.

Altogether, this computational finding strongly suggests that the strategically positioned chlorine atom effectively restricts the molecule’s conformational dynamics, likely enriching the population of a bioactive conformation pre-organized for productive ternary complex assembly. Such pre-organization is expected to minimize the entropic penalty of binding, thereby enhancing ternary complex formation and degradation efficacy, consistent with the experimental results. Therefore, reducing conformational flexibility represents a viable strategy for optimizing degrader efficacy.

### NGT-201-18 Induced VAV1 Degradation Inhibits Human T Cell Activation

To validate the functional consequences of VAV1 degradation in a physiologically relevant system, we evaluated the efficacy of the optimized degrader, NGT-201-18, in primary human T cells. VAV1 is a critical signal transducer downstream of the T-cell receptor (TCR), and its depletion is expected to blunt T cell activation. Treatment of primary T cells with NGT-201-18 resulted in robust, dose-dependent degradation of endogenous VAV1 (**Supplementary Figure 6A**). Following stimulation with anti-CD3/CD28 antibodies, this loss of VAV1 protein correlated with an impairment in T cell activation, as evidenced by the dose-dependent suppression of the cell surface activation marker CD69 (**Supplementary Figure 6B, C**). These results confirm that NGT-201-18 is cell-permeable and potent in primary human immune cells, effectively uncoupling TCR engagement from downstream activation signaling.

### Structure-Activity Relationship Studies on Phenyl Ring Substitutions Reveal Steric Tolerances

To further optimize the degrader scaffold and delineate the chemical space surrounding the core structure, comprehensive structure-activity relationship (SAR) studies were undertaken involving the introduction of diverse substituents at the ortho, meta, and para positions of the phenyl ring linking the phenyl-glutarimide based on NGT-201-17 (**Figure 5C**). This systematic investigation revealed distinct SAR trends. First, introduction of substituents at the ortho position relative to the phenyl ring consistently proves detrimental to VAV1 degradation activity (e.g., NGT-201-122, −124, −127, −133, −136, −139, −142, −145, −148). This finding indicates steric sensitivity at this position, potentially resulting from clashes with CRBN residues within the ternary complex. This correlates well with the ternary model that the ortho-position (4,6-indole position of NGT-201-18) is tightly flanked by CRBN H353 and W386 side chains (**Figure 5A, B**). In contrast, modifications at the para positions are generally well tolerated, with many substituents affording potent VAV1 degraders. This suggests greater solvent accessibility or available volume in these regions of the ternary complex interface. For example, analogs incorporating small para-NH_2_ or -OH (NGT-201-110/111) and para-CH_2_OH (NGT-201-146), or bulky para-NHCOCH_3_ (NGT-201-135), para-NHCONH_2_ (NGT-201-138), para-NHSO_2_CH_3_ (NGT-201-141), all exhibit favorable logDC_50_ values (superior to −7.5 of NGT-201-17), indicating tolerance for chemically diverse functional groups at these locations. Small polar moieties capable as hydrogen bond donors, including amine (-NH_2_ or -CH_2_NH_2_), hydroxyl (-OH or -CH_2_OH), are tolerated at the meta positions, but show to improve the potency when at the para position. Various hydrogen bond donor and acceptor-containing substituents including amide (NGT-201-135/138) and sulfonamide (NGT-201-141) also yield potent compounds only when situated at the para position. Overall, our SAR studies centered on the phenyl ring show strong concordance between experimental degradation data and predictions derived from the ternary complex model.

**Figure 5.**
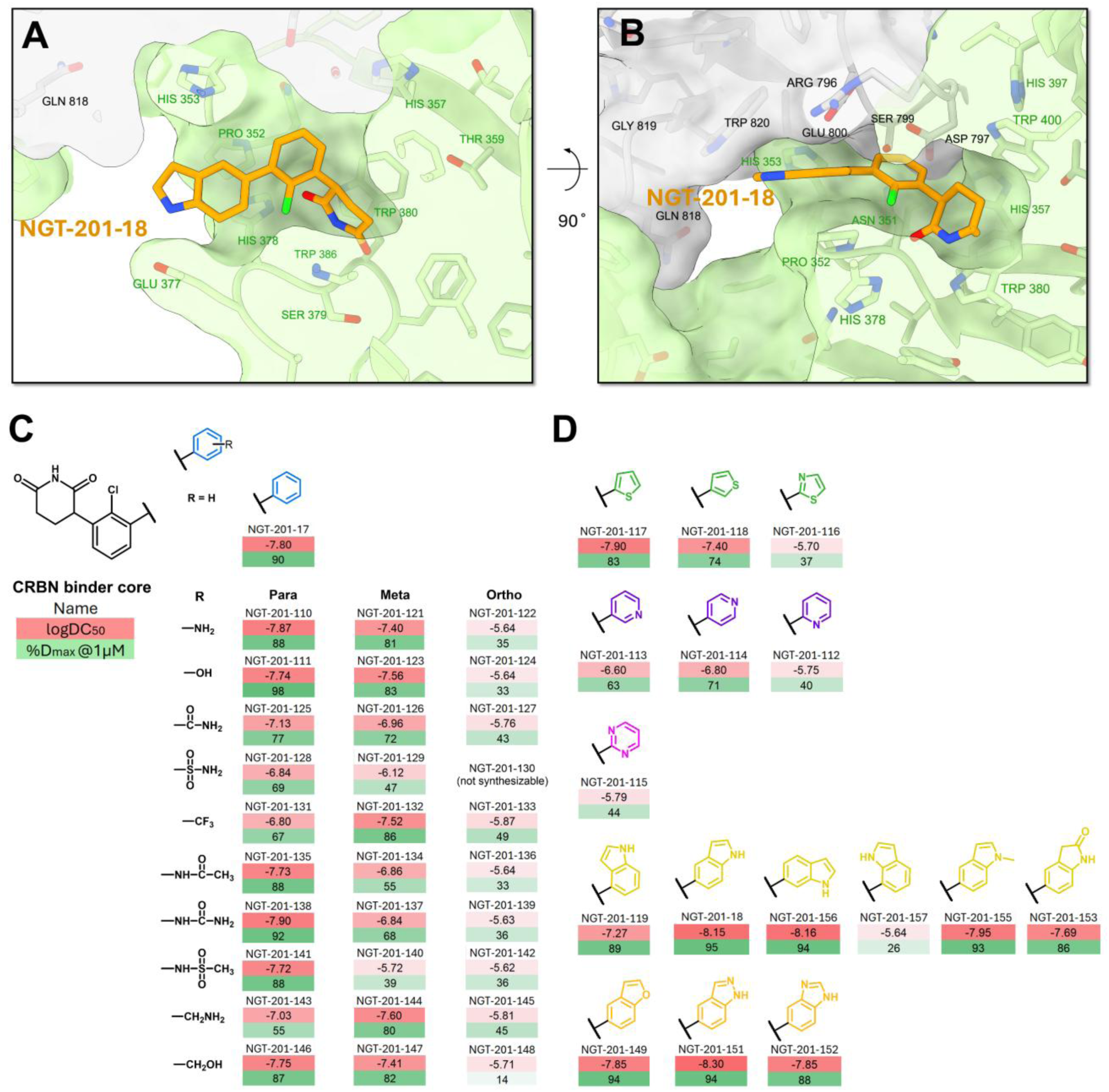
Structure-Activity Relationship (SAR) Studies on the Indole and Phenyl Moieties. **(A, B)** Two different perspectives of the computational model showing NGT-201-18 (orange sticks) bound within the pocket flanked by VAV1 SH3-2 (grey surface and sticks) and CRBN (light green). **(C)** SAR table summarizing the effect of various substituents at the ortho, meta, and para positions of the phenyl ring in the NGT-201 series on VAV1 degradation potency (logDC_50_) and efficacy (D_max_ at 1 μM). The core structure and the unsubstituted analog are shown at the top. Color coding indicates potency (green for higher %D_max_, red for stronger potency or lower DC_50_). **(D)** SAR table illustrating the impact of replacing the phenyl moiety of NGT-201-17 with different aromatic heterocycles on VAV1 degradation activity (logDC_50_) and efficacy (D_max_ at 1 μM). Color coding indicates potency (green for higher %D_max_, red for stronger potency or lower DC_50_).

### Structure-Activity Relationship Studies on the Indole Moiety Define Electronic Preferences

Following the identification of NGT-201-18 as a more potent VAV1 degrader possessing a conformationally restricted core and an indole VAV1-interacting element, the contribution of the indole ring itself was investigated. The indole ring compound by itself (NGT-201-18) is a more potent degrader than its phenyl counterpart (NGT-201-17). Analogs were synthesized and evaluated in which the indole moiety is substituted with alternative aromatic heterocycles (NGT-201-112 to −118, **Figure 5D**). Replacement of the electron-rich indole with comparatively electron-deficient heterocycles, such as pyridine (NGT-201-113, NGT-201-114) and thiazole (NGT-201-116), results in a reduction in VAV1 degradation potency. Replacement with pyrimidine (NGT-201-115), a more electron-deficient ring, leads to even more pronounced reduction in degradation potency (**Figure 5D**). Substitution with other phenyl bioisosteric heterocycles, particularly thiophene (NGT-201-117/118), yield compounds that maintained comparable activity. Substitution with 4-indole (NGT-201-119) or 7-indole (NGT-201-157) leads to reduced degradation potency, echoing the detrimental ortho-substitution discussed in the previous section. The nitrogen in the indole ring in NGT-201-12 appears to be a hydrogen bond donor to interact with E377 in CRBN. Compared with NGT-201-18, introducing a methyl group to the indole nitrogen as in NGT-201-155 only reduces its potency by < 3-fold, suggesting that the interaction with E377 in CRBN is dispensable. Replacing the indole ring in NGT-201-18 with benzofuran (NGT-201-149), benzopyrazole (NGT-201-151) and benzoimidazole (NGT-201-152) maintains the VAV1 degradation potency. All these observations suggest that the electron-rich character, and potentially the specific steric and electronic interaction profile (e.g., propensity for cation-π interaction suggested by modeling), of the indole and thiophene rings are favorable for productive engagement with the VAV1 SH3-2 domain within the ternary complex. Overall, all the congruence between the experimental SAR findings and the theoretical model predictions lends substantial support to the model’s accuracy and underscores its utility in guiding rational degrader design.

### FEP-based Calculation of Cooperativity Correlates with Experimental VAV1 Degradation Potency

To evaluate whether computational methods could predictively rank the degradation potency of synthesized VAV1 molecular glue analogs, we benchmarked multiple free-energy-based and docking-based scoring approaches against experimentally determined logDC_50_ and D_max_ values. FEP-based methods showed the strongest correlations with experimental potency, particularly for logDC_50_. Schrödinger FEP+ ternary complex calculations correlated strongly with logDC_50_ (Pearson r = 0.836, p = 0.0026), while OpenFE ternary calculations and AQFEP also showed significant correlations with logDC_50_ (Pearson r = 0.737, p = 0.015; Pearson r = 0.7856, p = 0.007, respectively) (**Figure 6A, C**). FEP-based cooperativity calculations also showed a significant inverse correlation with D_max_ (Pearson r = −0.655, p = 0.039) (**Figure 6A, C**), consistent with the expectation that more favorable interaction energetics are associated with improved degradation efficacy. In contrast, conventional docking-based methods, including Glide docking and Vinardo, generally showed weak or nonsignificant correlations with degradation potency, although CNNaffinity showed a modest correlation with D_max_ (Pearson r = −0.663, p = 0.037) (**Figure 6B, C**). These results support the use of free-energy-based calculations, rather than simple docking scores, for prospectively ranking and prioritizing VAV1 molecular glue analogs during lead optimization.

**Figure 6.**
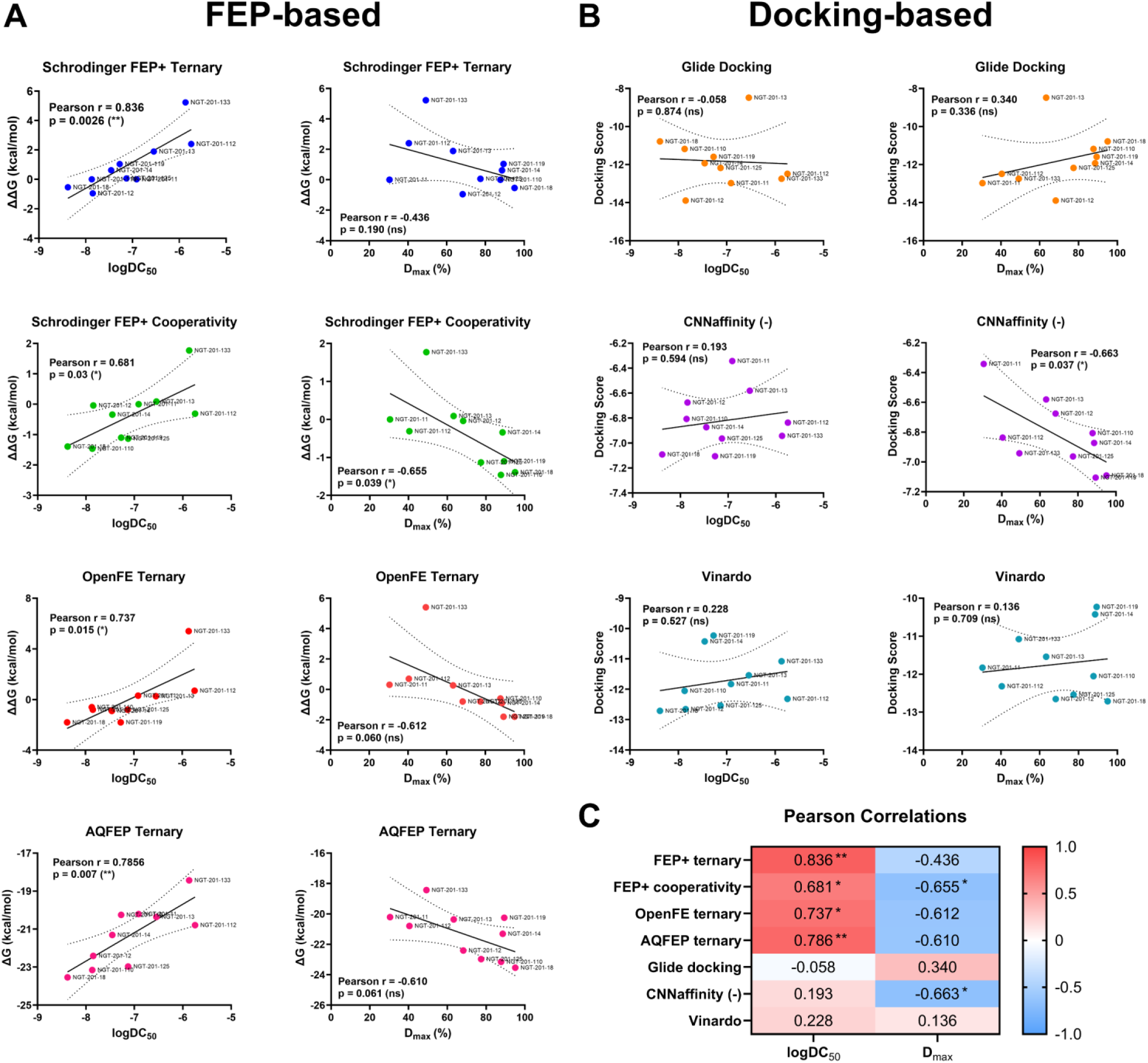
**Benchmarking FEP-based and docking-based computational metrics for predicting VAV1 degrader potency.** Correlation analyses comparing computationally calculated free-energy or docking-score metrics with experimentally determined VAV1 degradation potency across NGT analogs. (**A**) FEP-based approaches, including Schrödinger FEP+ ternary complex calculations, Schrödinger FEP+ cooperativity calculations, OpenFE ternary calculations, and AQFEP calculations, were compared against logDC_50_ and D_max_ at 1 μM. (**B**) docking-based approaches, including Glide Docking, CNNaffinity (docking score was inversed), and Vinardo, were compared against the same experimental endpoints. Each point represents one compound, with compound IDs indicated. Solid lines show the best-fit line from simple linear regression (the measure of centre), and dotted lines indicate its 95% confidence interval (n = 10 compounds). (**C**) Pearson correlation coefficients and corresponding significance from A and B.

With the FEP-based method accuracy confirmed, we next investigated the stereochemical preference of our degraders. Unlike standard lenalidomide derivatives that racemize rapidly, phenyl-glutarimide analogs exhibit slower racemization rates, rendering their stereochemistry a critical determinant for persistent neosubstrate engagement ^32,41^. We performed thermodynamic calculations using OpenFE (an open-source FEP approach) to compute the binding cooperativity of both the (R) and (S) enantiomers for NGT-201-11 and the conformationally restricted NGT-201-17 (**Supplementary Table 1**). The calculations revealed a distinct thermodynamic preference for the (R)-enantiomer during ternary complex formation. The highly potent chlorinated analog NGT-201-17(R) exhibited a strongly favorable cooperativity (ΔΔG_ternary_ = −1.14 kcal/mol, ΔΔG_coop_ = −1.52 kcal/mol), whereas its corresponding (S)-enantiomer was thermodynamically less favorable (ΔΔG_ternary_ = −0.43 kcal/mol, ΔΔG_coop_ = −0.88 kcal/mol). A parallel trend was observed for the unsubstituted NGT-201-11, where the (R)-enantiomer outperformed the (S)-isomer. These thermodynamic insights corroborate that the (R)-configuration, which structurally aligns with our GluePlex model and the canonical IMiD binding pose within the CRBN pocket ^32^, provides the optimal geometric and energetic cooperativity required for productive VAV1 neosubstrate recruitment.

### Dose-Response Proteomics Identifies LIMD1 as an Off-Target and Underscores the Utility of Proteomics

A fundamental requirement in the development of targeted therapeutics is the rigorous assessment of selectivity. To comprehensively evaluate the cellular target degradation profiles of the optimized VAV1 degraders, dose-response global proteomics analyses were conducted in Jurkat cells for several VAV1 degraders whose DC_50_ values are lower than 100 nM, including the highly potent NGT-201-18 (**Figure 7A, Supplementary Figure 7-8**). These experiments corroborated VAV1 as the primary target, exhibiting efficient degradation at low nanomolar concentrations. However, we noticed NGT-201-18 and many other compounds also induced dose dependent degradation of LIM Domain Containing 1 (LIMD1) protein. We further validated this finding orthogonally using a HiBiT-based reporter assay, which confirmed the degradation of transfected LIMD1 in response to NGT-201-18 treatment (**Figure 7B**).

**Figure 7.**
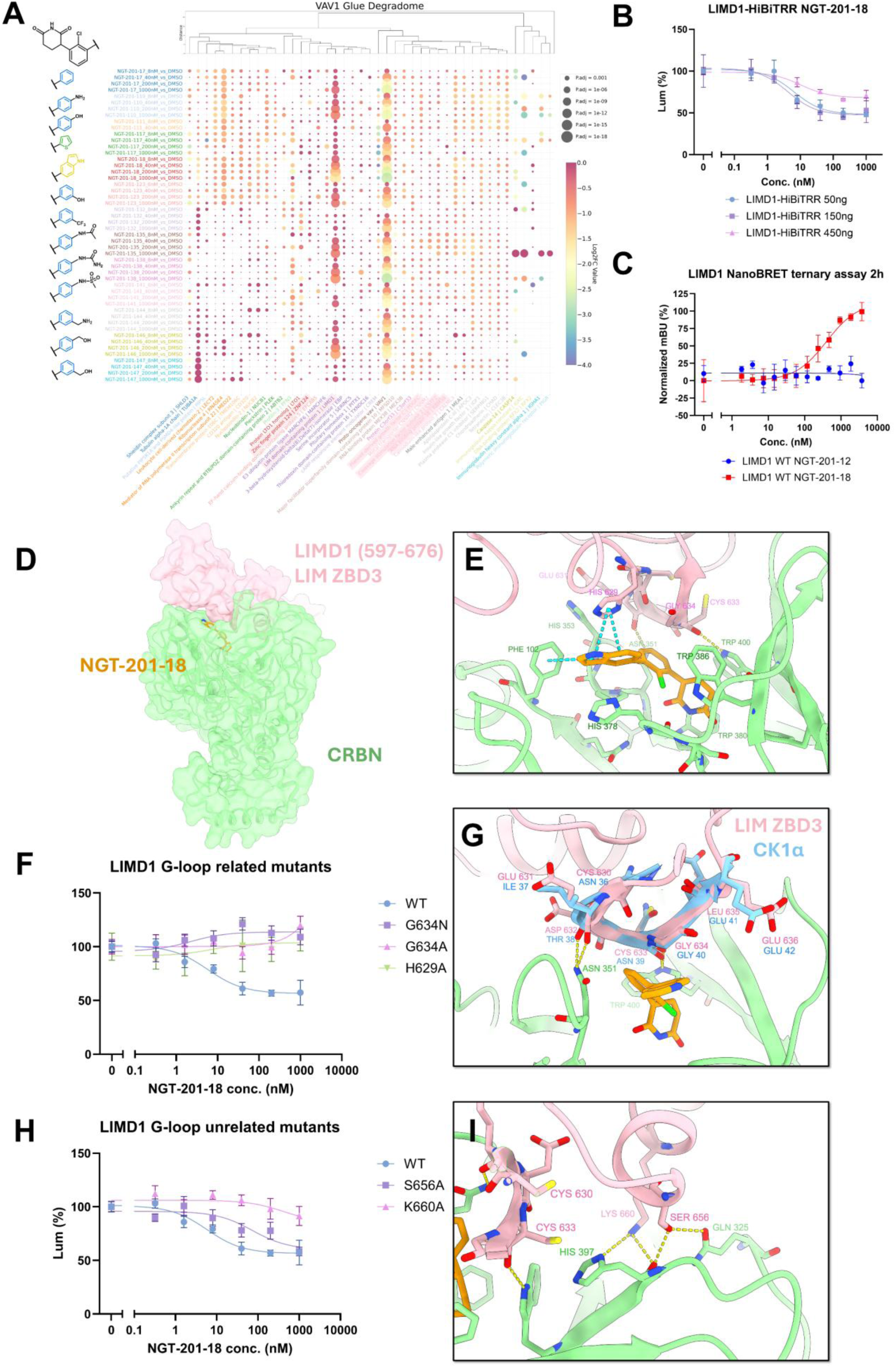
Dose-Response Proteomics Reveals LIMD1 as an Off-Target of VAV1 Glues. **(A)** Heatmap visualizing the dose-dependent changes in protein abundance for a panel of NGT compounds, highlighting VAV1 as the primary target in Jurkat cells. LIMD1 is also identified as a compound-dependent degradation target. The color intensity and circle size represent the magnitude and statistical significance of protein downregulation, respectively. Chemical structures of selected NGT compounds are shown on the left. Proteins harboring canonical G-loop degrons are highlighted in pink with their names. **(B)** HiBiT-based dose-response curves illustrating the degradation of transfected full-length LIMD1-HiBiTRR (plasmid 50 ng, 150 ng, 450 ng) upon 24 hours treatment with increasing concentrations of NGT-201-18 in Hela cells. Data are presented as mean±SD from three independent experiments. **(C)** Lysate-based NanoBRET LIMD1:degrader:CRBN ternary complex assay with nLuc-LIMD1 full length for NGT-201-12 / −18. Data are presented as mean ± SD from three independent experiments. **(D)** G-loop grafted structural model of the predicted ternary complex formed by CRBN (light green), NGT-201-18 (orange), and the LIMD1 zinc binding domain 3 (ZBD3, pink). **(E)** Close-up view of the predicted interaction interface between NGT-201-18 and LIMD1 ZBD3 domain, highlighting potential π-π interaction (cyan dashed lines) and hydrogen bonds (yellow dashed lines). **(F, H)** LIMD1-HiBiTRR WT or mutants were transfected into Hela cells and treated with various concentrations of NGT-201-18. H629 and G634 (F) are in the G-loop motif, while S656 and K660 (H) are at the PPI surface outside the G-loop motif. Data are presented as mean ± SD from three independent experiments. **(G)** Superimposition of the predicted NGT-201-18-bound LIMD1 ZBD3 structure (pink) with the known lenalidomide-bound CK1α G-loop structure (light blue), illustrating a similar binding mode within the CRBN pocket (green). **(I)** Close-up view of the predicted G-loop unrelated interaction interface between CRBN and LIMD1 ZBD3 domain, highlighting potential hydrogen bonds (yellow dashed lines).

To confirm that LIMD1 degradation occurs via a molecular glue mechanism, we performed a NanoBRET-based ternary complex assay. The results demonstrated that NGT-201-18, but not NGT-201-12, effectively promotes the formation of a ternary complex between CRBN and LIMD1 (**Figure 7C**). Notably, sequence and structural analysis revealed that LIMD1 harbors a sequence motif consistent with the canonical G-loop degron (residue 630-636) located in the Zinc Binding Domain 3 (ZBD3) (**Figure 7G**). Structural modeling of the CRBN:NGT-201-18:LIMD1 ternary complex predicted that the indole moiety of NGT-201-18 engages in a π-π stacking interaction with LIMD1 His629, a key residue within the G-loop (**Figure 7D-E**, **Supplementary Figure 9**, **Supplementary Data 2**). Similar with CK1*α* G-loop degron, our model highlighted the potential hydrogen bonds between CRBN and LIMD1 D632 and C633 backbone carbonyls (**Figure 7G**). This structural hypothesis aligns with our SAR data, which shows that degraders with electron-rich rings like indole exhibit stronger degradation for both VAV1 and LIMD1, while compounds with benzene or thiophene rings spares LIMD1 from degradation (**Figure 7A, Supplementary Figure 7-8**).

To functionally validate this predicted interface, we performed site-directed mutagenesis on LIMD1. As hypothesized, mutating key sidechain interacting residues within the G-loop degron (H629A, G634N or G634A) completely abrogated NGT-201-18-induced LIMD1 degradation (**Figure 7F**). Furthermore, mutating residues predicted to be on the broader protein-protein interface but outside the G-loop, such as S656A and K660A, also impaired LIMD1 degradation (**Figure 7H, I**). These data collectively confirm that NGT-201-18 recruits LIMD1 to CRBN through a canonical G-loop-dependent mechanism.

This significant finding illustrates that a single molecular glue compound may possess the capacity to recognize and mediate the degradation of neosubstrates bearing distinct degron motifs (the non-canonical RT loop in VAV1 and a canonical G-loop in LIMD1). This powerfully underscores the indispensable value of systematic, unbiased global proteomics throughout the molecular glue discovery and optimization process. Such methodologies are crucial for obtaining a comprehensive understanding of both on-target and potential off-target activities, which are critical for developing safe and effective degraders.

### GluePlex is Essential for Predicting Non-Canonical Degron Interfaces

To explicitly validate the necessity of the integrated GluePlex workflow for de novo degron discovery, we benchmarked its performance against the isolated Boltz-2 co-folding algorithm using both our canonical (LIMD1) and non-canonical (VAV1) targets. When the full GluePlex pipeline was applied to LIMD1, almost all HADDOCK docking templates successfully converged into the correct G-loop binding pose during co-folding, as demonstrated by the PCA mapping of the generated structures (**Supplementary Figure 10A**). Furthermore, when tasked with predicting the LIMD1:NGT-201-18:CRBN ternary complex without any prior HADDOCK docking templates (with MSA generation explicitly disabled, but provided with individual proteins as templates), the standalone Boltz-2 model successfully generated high-confidence structures with low RMSD relative to our alignment-based reference (**Supplementary Figure 10B**). This robust performance is anticipated, as LIMD1 utilizes a canonical G-loop degron whose structural motif is highly enriched in the training datasets of AlphaFold-derived models.

In contrast, when the isolated Boltz-2 algorithm was applied to the non-canonical VAV1 RT-loop degron, it completely failed to generate viable ternary assemblies. The standalone model yielded exclusively low confidence scores (chain-pair ipTM < 0.3) and extreme structural deviations (Cα RMSD > 10 Å) (**Supplementary Figure 10C**). This comparative analysis firmly establishes that while current standalone deep learning folding models can successfully recall known G-loop paradigms due to training data bias, the integration of PeSTo and HADDOCK to provide physics-based structural restraints within the GluePlex pipeline is strictly indispensable for resolving non-canonical neosubstrate interfaces.

### Proteome-Wide Search Suggests Potential RT Loop Mimics

The characterization of the functional, non-canonical RDxS motif within the VAV1 SH3-2 RT loop as a CRBN-dependent degron motivated an investigation into its potential occurrence elsewhere within the human proteome. Bioinformatic research was conducted to identify proteins containing similar sequences, with particular emphasis on those presented within structurally analogous contexts, such as the RT loops of other SH3 domains. Sequence alignments coupled with structural comparisons identified several candidate proteins possessing SH3 domains with RT loop sequences exhibiting similarity to the VAV1 degron (**Supplementary Figure 11-12**). Prominent examples include the GRB2-related adaptor protein, as well as the Endophilin-B2. These proteins constitute theoretical off-targets requiring consideration during the selectivity assessment of VAV1 degraders. From the other perspective, this compilation also furnishes a set of potential starting points for the rational design of molecular glues intended to degrade these specific targets by exploiting the structural principles elucidated in the VAV1 system.

## Discussion

Our study successfully identified and characterized a series of CRBN molecular glues that selectively target the hematopoietic-specific signaling protein VAV1 for proteasomal degradation. Utilizing an unbiased, deep global proteomics approach, we discovered that phenyl-glutarimide derivatives, notably NGT-201-12, can effectively induce VAV1 degradation. Mechanistic investigations pinpointed the C-terminal SH3 domain (SH3-2) of VAV1 as crucial for this process. This research expands our understanding of molecular glue mechanisms and offers a strategy for targeting VAV1, a protein implicated in various hematological malignancies and autoimmune diseases.

A key finding of this work is the elucidation of a non-canonical degron within the VAV1 SH3-2 domain. While CRBN molecular glues are often associated with the recognition of G-loop degrons, our computational modeling and subsequent site-directed mutagenesis studies compellingly demonstrated that an RT loop, specifically an RDxS motif (residues 796-799), is essential for the NGT-201-12-mediated degradation of VAV1. Consistent with our finding, the recent independent report by Petzold et al. ^19^ highlights the versatility of CRBN as an E3 ligase capable of recognizing diverse structural motifs on neosubstrates, moving beyond the canonical G-loop paradigm. The agreement between two studies using orthogonal approaches provides mutual support for assignment of the SH3-2 RT loop as the recognition element. The identification and characterization of this non-canonical RT-loop degron provides a basis for designing molecular glues against targets previously considered intractable due to the absence of known degron motifs.

This study further underscores the potent synergy between predictive structural modeling and iterative chemical optimization in the development of effective protein degraders. We developed the GluePlex workflow, a physics-and AI-driven modeling tool prioritizing protein-protein interactions with small molecule ligands. This platform successfully predicted the key intermolecular contacts within the ternary complex formed by our lead compound, CRBN, and the VAV1 SH3-2 domain, including crucial PPIs between CRBN and VAV1. These structural hypotheses were subsequently validated experimentally through our site-directed mutagenesis studies. Our predicted VAV1 ternary complex structure aligns closely with the crystal structure of CRBN-DDB1 and MRT-23227 in complex with VAV1 (PDB: 9NFR) as reported by Petzold et al. ^19^. Notably, the Boltz-2 model was released prior to the 9NFR structure released in the Protein Data Bank, ruling out the possibility that the model “memorized” the crystal structure and confirming the predictive power of the workflow. Retrospectively, we evaluated the structural fidelity of the input templates relative to the final predicted ternary complex. Even the closest protein-protein docking model (HADDOCK Cluster 4) exhibited an RMSD of approximately 8 Å for the VAV1 SH3-2 domain compared to the final co-folded structure. Despite this significant initial deviation, the Boltz-2 algorithm successfully converged to a high-confidence binding pose (ipTM = 0.94) during the co-folding stage. This indicates that the generative model is not merely rigid-body fitting the ligand into a static pocket but possesses substantial capacity for structural refinement. The ability to traverse from a coarse-grained protein-protein docking template to a precise atomic-resolution ternary interface suggests that GluePlex maintains high generalizability and can effectively locate the local energetic minimum even when provided with imperfect starting geometries.

While techniques like Cryo-EM and X-ray crystallography remain the gold standard for high-resolution structural elucidation, their demanding requirements for highly optimized compounds and experimental conditions often mean that ternary complex structures are determined retrospectively. As a result, their direct impact on prospectively guiding the initial hit-to-lead optimization in drug discovery can be limited, with many reported structures serving as post-hoc analyses. In contrast, the GluePlex workflow offers a significant strategic advantage by enabling the generation of actionable structural models of ternary complexes during the critical early phases of discovery. This study demonstrates that when coupled with rigorous experimental validation such as the site-directed mutagenesis and SAR reported here, these predictive models can prospectively guide medicinal chemistry. This approach empowers researchers to transition from unbiased proteomic hits to structure-based drug design strategies, facilitating the efficient engineering of compounds with enhanced potency and selectivity before high-resolution experimental structures are available.

Additionally, our ternary complex structural modeling suggested that the conformational flexibility in NGT-201-12 could be a liability. Strategic chemical modifications, particularly conformational restriction through halogen substitution (e.g., NGT-201-17, NGT-201-18), led to potentiatied VAV1 degradation. Density Functional Theory calculations supported the hypothesis that these modifications reduce conformational flexibility, thereby minimizing the entropic penalty associated with ternary complex formation and enhancing degradation efficacy. This principle of molecular rigidification offers a valuable strategy for optimizing the potency of other molecular glue degraders.

The comprehensive structure-activity relationship (SAR) studies provided further insights into the chemical features governing degrader activities. These studies delineated specific steric tolerances and electronic preferences for substituents on both the phenyl and indole moieties of the degrader scaffold. For instance, substitutions at the ortho position of the indole-linking phenyl ring were generally detrimental, correlating with predicted steric clashes within the ternary complex model. Conversely, meta and para positions showed greater tolerance for diverse chemical groups. Investigations into the indole moiety itself revealed a preference for electron-rich aromatic heterocycles, likely contributing to favorable interactions, such as cation-π interactions, with the VAV1 SH3-2 domain. These detailed SAR findings provide a roadmap for the rational design of next-generation VAV1 degraders with improved properties.

A critical aspect highlighted by our research is the indispensable role of unbiased, global proteomics in the discovery and optimization of molecular glues. Enabled by SP3-based high throughput sample preparation ^42,43^, Orbitrap Astral MS platform and downstream DIA data analysis workflow ^44^, dose-response proteomics analyses not only confirmed VAV1 as the primary target of our optimized compounds but also unexpectedly revealed LIMD1 as an off-target for some analogs, including the potent NGT-201-18. LIMD1 possesses a canonical G-loop-like degron, and structural modeling supported the ability of NGT-201-18 to facilitate CRBN interaction with this motif. This demonstrates that a single molecular glue can engage distinct degron motifs on different neosubstrates, a finding with significant implications for selectivity profiling and drug development. It underscores the necessity of comprehensive proteomic assessment to understand both on-target and potential off-target liabilities, guiding the development of safer and more effective molecular glue degraders.

For the LIMD1 ternary complex modeling, we deliberately employed a scientifically robust and efficient structural alignment strategy, leveraging the hypothesis that LIMD1 utilizes a canonical G-loop degron. This well-characterized motif has a known binding mode with CRBN, allowing us to generate a high-confidence model by directly aligning the LIMD1 G-loop with the solved crystal structure of the CK1α:lenalidomide:CRBN complex (PDB: 5FQD). The accuracy of our alignment-based LIMD1 model was subsequently confirmed by our mutagenesis studies, validating this judicious choice of a more traditional and efficient modeling strategy. We further retrospectively evaluated the standalone co-folding algorithm (Boltz2 with no MSA) on both proteins. While the standalone Boltz-2 model successfully generated high-confidence predictions for the canonical LIMD1 complex—likely due to the enrichment of G-loop motifs in its training data—it completely failed to assemble the unprecedented VAV1 RT-loop interface, yielding severe structural deviations and low confidence scores. This contrast highlights that while current deep learning models can efficiently recall established degron paradigms, integrating physics-based structural restraints, as done in GluePlex, remains strictly indispensable for predicting entirely novel, non-canonical neosubstrate interactions.

Finally, the identification of the functional RDxS motif in VAV1 prompted a proteome-wide search for similar sequences, particularly within structurally analogous RT loops of other SH3 domain-containing proteins. This bioinformatic analysis identified several potential candidates, such as GRB2-related adaptor protein (GRAP) and Endophilin-B2. These proteins, on one hand, represent potential off-targets in non-targeted tissues that warrant consideration during further development of VAV1 degraders. Conversely, this knowledge also provides a foundation for the rational design of molecular glues aimed at degrading these specific targets by leveraging the structural principles uncovered in the VAV1 system.

In conclusion, this study has unveiled a non-canonical RT-loop degron in VAV1 that is recognized by CRBN molecular glues, demonstrated the utility of conformational restriction in potentiating degrader activity, and highlighted the crucial role of combining global proteomics and advanced modeling techniques in understanding degrader potency and selectivity. These findings not only provide a therapeutic strategy for targeting VAV1 but also contribute to the broader field of targeted protein degradation by expanding the known landscape of CRBN neosubstrate recognition and informing the rational design of molecular glue degraders. Future work will focus on further optimizing the selectivity and in vivo efficacy of these VAV1 degraders and exploring the potential for targeting other proteins via similar RT-loop degrons.

## Methods

### Ethics statement

This study did not involve the recruitment of human participants by the investigators. Primary human peripheral blood mononuclear cells (PBMCs) were obtained as de-identified leukocyte units from healthy adult volunteer donors through the Gulf Coast Regional Blood Center (Houston, TX, USA). Written informed consent was obtained from all donors by the Gulf Coast Regional Blood Center prior to sample collection, including consent for the use of donated blood products in research. No donor is an author of this study, and the investigators had no access to identifiable private information at any point. Accordingly, the use of these samples was reviewed by the Institutional Review Board for Human Subject Research at Baylor College of Medicine, which determined that this work does not constitute human subjects research as defined by 45 CFR 46.102(e) and issued an exemption/waiver (determination no. [H-39503]). All experiments involving human-derived material were performed in accordance with the ethical principles of the Declaration of Helsinki.

### Cell culture & Transfection

Jurkat (ATCC, TIB-152), HEK293T/17 (ATCC, CRL-11268), and HeLa (ATCC, CCL-2) cells were purchased from ATCC. Jurkat cells were cultured in RPMI-1640 (Corning, 10-040-CV) supplemented with 10% FBS (Corning, 35-011-CV). HEK293T and Hela cells were cultured in DMEM (Corning, 10-013-CV) supplemented with 10% FBS. All the cell cultures are in the 37 °C incubator with 5% CO_2_. Opti-MEM Reduced Serum Medium used for transfection is from Gibco 11058021.

For degradation assay, nLuc-VAV1 plasmids with truncations or mutations were transfected into Hela cells using Lipomaster 3000 (Vazyme, TL301-01). Hela cells (0.3 million per well) were plated in the tissue culture treated 6-well plate the day before transfection. Immediately before transfection, the media were refreshed. Liposome transfection reagents were prepared by mixing tube A (Opti-MEM 125 μL + Lipomaster 3000 3.75uL) and B (Opti-MEM 125 μL + DNA 150 ng +T3000 enhancer 4 μL), waiting for 10min, and dropwise added into the culture medium. After 24h expression, cells were harvested for the degradation assay.

For CRBN NanoBRET target engagement assay, nLuc-CRBN and DDB1 plasmids (Promega, N2741 and N2761) were transfected into HEK293T cells using calcium phosphate method. Briefly, 2.5 million HEK293T cells in 10mL complete medium were plated in 10cm petri dishes and allowed for 4h settlement. Calcium phosphate-DNA particles were formed by mixing 1 μg nLuc-CRBN plasmid, 4 μg DDB1 plasmid, 75 μL 2M CaCl_2_, 540 μL ddH2O and 625 μL 2x HBSS, and incubate for 15min RT, then dropwise added into the petri dish. After 18h, the media were refreshed and cells were harvested after another 24h for NanoBRET assay.

### VAV1 HiBiT-RR CRISPR knock-in

Cells were subcultured three days before electroporation. The culture medium was replaced the day before the procedure. CRISPR-Cas9 sgRNAs and HDR ssODN templates were resuspended in IDT buffer E. sgRNA (CGTCTCTCTGCCA<u>AGG</u>CACC) was prepared at a final concentration of 100 µM by dissolving 2 nmol in 20 µL of buffer. Cas9 protein (62 µM in 50% glycerol, Integrated DNA Technologies Inc, 1081058) was diluted to 40 µM using Resuspension Buffer R from the Neon Transfection Kit (Invitrogen, MPK10025). For each reaction, 0.5 µL of 100 µM sgRNA was combined with 0.5 µL of 40 µM Cas9 in a total volume of 1.0 μL. The mixture was incubated at room temperature for 15 minutes to form the ribonucleoprotein (RNP) complex. Complete growth media without antibiotics was pre-warmed in a 37°C incubator, supplemented with a 500× dilution of Alt-R™ HDR Enhancer V2 (Integrated DNA Technologies Inc, 10007910). Jurkat cells were washed with 5 mL PBS to remove FBS and resuspended in Buffer R at a concentration of approximately 2 × 10⁷ cells/mL. For each electroporation, 8 μL of cell suspension (approximately 1.6 × 10⁵ cells) was used. In a 200 μL PCR tube, 1 μL of the RNP complex, 8 μL of the prepared cell suspension, and 1 μL of 100 µM HDR template (Alt-R HDR Donor Oligo, Integrated DNA Technologies Inc, Sequence: GCACTGATGAACTCCTCGTCTGTTTCCAGGTTGGCTGGTTCCCTGCCAACTACGTGGAGGAAGATTATTCTGAATACTGCGTCTCCGTGAGCGGCTGGCGGCTGTTCCGCAGGATTAGCTGAGCGGTGGTGCCTTGGCAGAGAGACGAGAAACTCCAGGCTCTGAGCCCGGCGTGGGCAGGCAGCGGAGCCAGGGGCTGT) were combined, bringing the total volume to 10 μL. The mixture was pipetted into a Neon Tip, ensuring no bubbles were present. Electroporation was performed using the Neon Transfection System with the following parameters: 1400 V, 30 ms pulse width, and 2 pulses.

Following electroporation, cells were immediately transferred to pre-warmed antibiotic-free media containing 500× diluted HDR enhancer. After 24 hours, the medium was replaced with standard culture medium. Cells were incubated for 72 hours at 37 °C before downstream analysis. Pooled HiBiT-RR knock-in Jurkat cells were used for high throughput VAV1 degradation assay.

### High-throughput global proteomics

#### SP3-based sample preparation

Wildtype Jurkat cells in complete media were plated in the 96-well U-bottom plate at a density of 200,000 cells per well. After 2 hours settlement, compounds dissolved in DMSO were micro-dispensed by Pico 8 to the corresponding concentration in each well and returned to the 37°C incubator for 24 hours treatment. Cells were centrifuged down, and the media was aspirated, followed by 1x PBS washing once. Cell pellets were lysed in 50 μL 200 mM HEPES, 0.25% SDS at pH 8.5, with 1x Halt protease inhibitor cocktail (Thermo Scientific, 78440) and 10U/well benzonase (Sigma, E1014) with 800 rpm shaking at 37°C for 30 min. Reduction was performed by adding 5 μL 50mM DTT and shaking at 37°C for 15 min, followed by alkylation with 14 μL 100 mM iodoacetamide at room temperature in dark for 30 min. To each well, 100 ug SP3 beads (1:1 mixture of E3 and E7 carboxylic magnetic beads from Cytiva) were added. Protein binding was initiated by adding 75 μL 100% ethanol into each well and shaking at 800 rpm for 10 min at room temperature. Beads were washed in the plate with the aid of magnetic rack using 200 μL 80% ethanol for twice. After the final wash, beads were briefly dried, and proteins were digested in 30 μL 12.5 ng/μL trypsin/LysC (Promega, V5073) in 200 mM HEPES with 1,000 rpm shaking at 37 °C for 18 hours, with the plate filmed.

#### Peptide clean-up with StageTip in G6-well plate format

The digested peptides were acidified with 1ul/well formic acid (FA) and further desalted and cleaned up with StageTip (CDS Analytical, 6091). The whole box of StageTip was transferred to a tip box which allows the placement of a 96-well non-skirted PCR plate inside for eluent collection purpose (Genesee Scientific, 22-102). Tips were moistened by adding 30 μL 80% ACN/0.1% FA, centrifuging the whole tip box at 125 x g for 1 min, and removing the liquid in the box. Plunge with air to remove excess liquid from the tip if necessary. Tips were equilibrated by adding 30 μL 0.1%FA and centrifuging and plunging as described above. The plated digests were transferred into 96 StageTips followed by centrifuging and plunging.

StageTips were washed twice with 30 μL 0.1% FA, and eluted with 100 μL 80% ACN/0.1% FA into a 96-well PCR collecting plate. The eluent plate were speed vac to dry, and peptides were resuspended in 30 μL 5% ACN/0.1% FA. Peptide concentration was measured with Fluorescent Quantitative Peptide Assays (Pierce, 23290) and normalized across the plate before LC-MS/MS injection.

#### Data-independent Acquisition and LC-MS/MS

All the LC-MS/MS data was collected on the Vanquish Neo coupled to Orbitrap Astral mass spectrometer through an EASY-Spray source (Thermo Fisher Scientifics). 300 ng – 1 μg peptides were loaded onto the 300μm×5mm trap column (Thermo Scientifics, 174500) and eluted onto the main 150um×15cm C18 column (Thermo Scientifics, ES906) using the Trap-and-Elute workflow. Peptides were eluted out to the spray with a 5-40% gradient of ACN/0.1% FA in 23 min. Nanospray ionization was operated at +2.0 kV, the ion transfer tube was held at 300 °C, the collision gas pressure was 1 mTorr, and total carrier gas flow was static. Neither the APPI lamp nor FAIMS interface was used. EASY-IC™ lock mass correction was enabled at RunStart, the application mode was set to “Peptide” under standard pressure conditions, the expected chromatographic peak width was defined as 15 s, the default precursor charge state was set to 2, and advanced peak determination was turned on. Full-scan (MS1) data were acquired from 0 to 23 min in the Orbitrap at a resolution of 240,000 (at m/z 200) across an m/z range of 380–980. The AGC target was set to 5 × 10^6^ (normalized AGC target 500%), with a maximum injection time of 5 ms, one microscan, and the RF lens set to 40%. Data were recorded in profile mode, in positive polarity, with source-induced fragmentation and lock-mass injection both disabled. Data-independent acquisition (MS2) was performed concurrently over the same 0–23 min window using the Astral detector. A total of 300 sequential isolation windows, each 2 Th wide and placed automatically without overlap, covered the precursor range of 380– 980 m/z. MS2 scans were collected in centroid mode across 150–2,000 m/z, with HCD normalized collision energy of 25%, an AGC target of 5 × 10^4^ (normalized AGC target 500%), maximum injection time of 5 ms, and one microscan. Cycle time was controlled by a 0.6 s loop in Start/End Time mode.

#### DIA data analysis

Raw files from Orbitrap Astral Mass Spectrometer were converted into the mzML format using ThermoRawFileParser1.4.5 and searched with DIA-NN 1.9.2 on the Linux platform ^44^. A spectral library was generated *in silico* from the provided human FASTA database and refined based on the DIA data (--gen-spec-lib, --predictor). The search was conducted against the reviewed UniProtKB *Homo sapiens* proteome (UP000005640, downloaded on Jan 7, 2025).

Search parameters included: a precursor m/z range of 380−980 and a fragment m/z range of 200−2000. Peptides between 6 and 40 amino acids in length with precursor charge states ranging from 2 to 6 were considered. Trypsin/P digestion rules were applied (--cut K*,R*), allowing for up to one missed cleavage. N-terminal methionine excision (--met-excision) was enabled. Mass accuracy was set to 15 ppm for MS1 precursors and 20 ppm for MS/MS fragments. Variable modifications included oxidation of methionine (UniMod:35, 15.994915 Da), with a maximum of two variable modifications per peptide. The --unimod4 flag was active for fixed carbamidomethylation of cysteine.

The analysis employed a double-search strategy (--double-search) and retention time-dependent scoring and profiling (--rt-profiling) for improved identification and quantification. Protein inference was performed using relaxed settings (--relaxed-prot-inf), and protein groups were reported (--pg-level 1). Quantification was based on extracted ion chromatograms (XICs) (--xic), and quantification method using QuantUMS ^45^ in high-precision mode. A false discovery rate (FDR) of 1% was applied at both peptide and protein levels (--qvalue 0.01).

Generated protein group quantification report, after filtering out the common contaminant proteins, was used to perform differential analysis and statistics using limma model on R 4.4.2 with Benjamini-Hochberg correction to control the False Discovery Rate (FDR).

### Immunoblotting

Wildtype Jurkat cells were seeded at 0.2 million cells per well in 1 mL of media in 24-well plates. After 24 hours of degrader treatment, cells were centrifuged, washed with PBS, and lysed in 1× RIPA buffer (Thermo Scientific, J62524.AE) supplemented with 1× Halt™ Protease and Phosphatase Inhibitor Cocktail (Thermo Fisher Scientific, Cat. No. PI78440) and Benzonase Nuclease (Sigma, E1014) to reduce viscosity. Lysates were cleared by centrifugation at 15,000 × g for 10 minutes at 4 °C, and total protein concentration was determined using the Pierce BCA Protein Assay Kit (Thermo Fisher Scientific, Cat. No. 23225). 20 µg of total protein were resolved by SDS-PAGE and transferred to a 0.2 µm PVDF membrane (Millipore, Cat. No. IPVH00010) pre-soaked in ethanol. Membranes were blocked and incubated overnight at 4 °C with primary antibodies diluted at 1:1000 (VAV1, Cell Signaling Technology #4657, clone D5G6, lot 2) or 1:4000 (β-actin, Cell Signaling Technology #4970, clone 13ES, lot 19), followed by three washes with 1× TBST. Blots were then incubated with HRP-conjugated secondary antibody (Kindle Biosciences, #R1006, 1:1000) for 1 hour at room temperature. After three additional washes with 1× TBST, detection was performed using 3× diluted SuperFemto ECL substrate (Vazyme, E423-01) and imaged using KwikQuant digital imaging system.

### Molecular cloning and mutagenesis

Wildtype human full length VAV1 gene fused with N-terminal nLuc was synthesized and cloned into pRP[Exp]-Puro-CMV by VectorBuilder Inc. Wildtype human full length LIMD1 cDNA was obtained from the Advanced Cell Engineering & 3D Models Core at Baylor College of Medicine and cloned into the same backbone by replacing VAV1. PCR amplicons were obtained using KOD One (Sigma, KMM-101NV) with specified mutagenesis PCR primer pairs (manufactured by Integrated DNA Technologies, Inc.) in the **Table 1**, followed by DpnI digestion (New England Biolabs, R0176S) and PCR product clean-up using FastPure gel extraction kit (Vazyme, DC301-01). Both site-directed mutagenesis and domain truncation were performed using ClonExpress Ultra One Step Cloning Kit V2 (Vazyme, C116-01) under 50 °C for 15 min. Homologous recombination products were directly transformed into DH5α competent cells (Thermo Scientific, EC0112) and colonies were grown under ampicillin selection pressure. Product plasmids were all validated through whole plasmid sequencing.

**Table 1.**
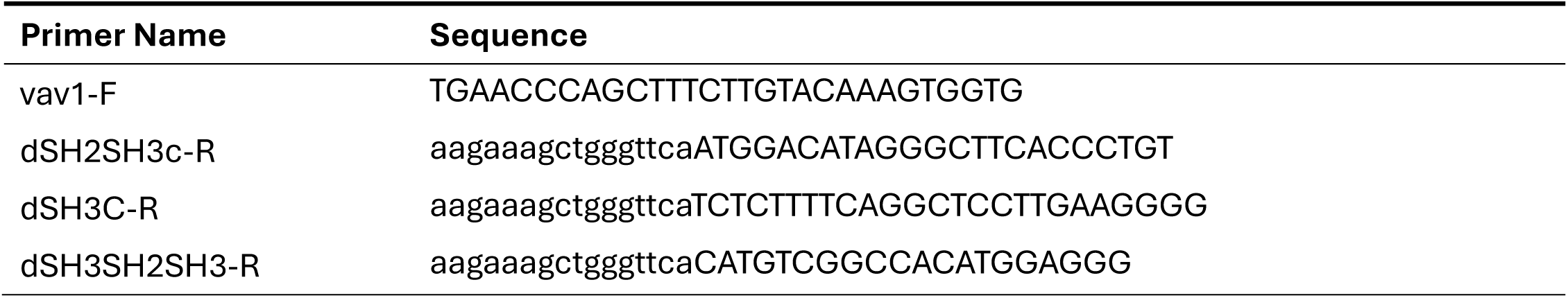
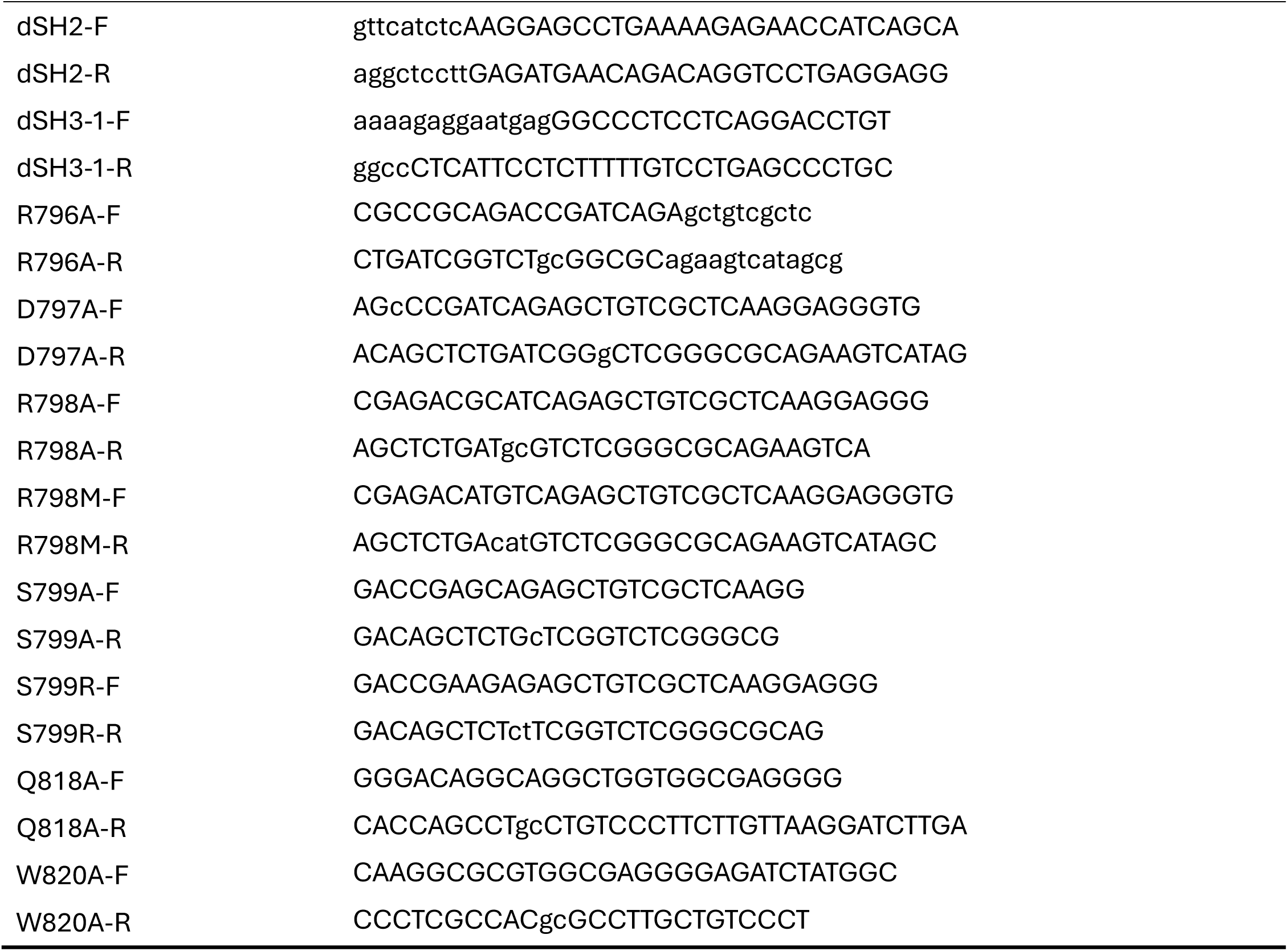
Primers used for VAV1 mutants cloning.

### HiBiT degradation assay of endogenous VAV1

Jurkat VAV1-HiBiT-RR knock-in cells were plated in the white 96-well plate (Corning, 3917) with a density of 20,000 cells/well. After overnight incubation, degrader molecules dissolved in DMSO were micro-dispensed with Multidrop™ Pico 8 Digital Dispenser (Thermo Fisher Scientific) to achieve 5-fold serial dilution in the culture medium. Cells were returned to the 37°C incubator for 24 hours incubation. After incubation, 100 μL NanoGlo cell lysis buffer containing 100-fold diluted NanoGlo substrate (Promega, N1573) and 200nM LgBiT (Promega, N401A) were added into each well. After brief shaking, luminescence signal was measured using H1 synergy (BioTek Instruments, Inc.) using a PMT gain of 120.

### Nano luciferase degradation assay of exogenous VAV1

Hela cells transfected with nLuc-VAV1 variants were trypsinized, centrifuged, and resuspended in the phenol red-free DMEM (Corning, 17-205) supplemented with 10% FBS. Cells were re-plated into white 96-well plates with a density of 20,000 cells/well in 100 μL. Degrader molecules in DMSO were micro-dispensed as described previously. After 24 hours treatment, 100 μL NanoGlo lysis buffer containing 200-fold diluted NanoGlo substrate were added into each well. After brief shaking, luminescence signal was measured using BioTek H1 synergy using a PMT gain of 120.

### NanoBRET-based CRBN target engagement assay

HEK293T cells transfected with nLuc-CRBN and DDB1 plasmids were trypsinized, centrifuged, and resuspended in the phenol red-free DMEM (Corning, 17-205) supplemented with 10% FBS and 500 nM CRBN NanoBRET tracer molecule (Promega, CS1810C137). At this concentration without any other competitor compounds, the tracer can achieve 50% maximal target engagement on CRBN using the exact same protocol described below. Cells with tracer-containing media were re-plated into white 96-well plate with a density of 40,000 cells/well in 100 μL and incubated for 1 hour. Degrader molecules with 5-fold dilution in DMSO were dispensed in equi-volume. After 2 hours target engagement, 25 μL of PBS containing 100-fold diluted NanoGlo substrate and 10 μM extracellular nLuc inhibitor (Promega, N235A) was added into each well. Dual channel signal intensities were simultaneously measured on a BMG PheraStar plate reader mounted with 450±80nm and 610nm LP filter set, with auto gain setting on both channels, in precision mode. BRET ratio in milli-BRET unit (mBU) were calculated using Equation 1:

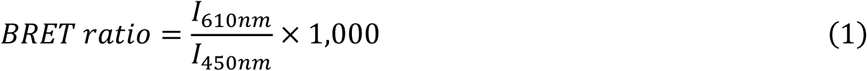

### NanoBRET-based ternary complex formation assay

About 1.2 million Hela cells transfected with nLuc-VAV1 or nLuc-LIMD1 WT or mutants were washed with PBS and lyzed in 500 μL NP40 lysis buffer (50 mM Tris-HCl pH 7.5, 250 mM NaCl, 0.5% NP-40 alternatives, 10% glycerol, 1 mM EDTA, 1x Halt Protease inhibitor cocktails). All 500 μL lysate were incubated with 500 μL 400 nM HisCRBN/HisDDB1 purified protein, 500 μL 800 nM HIS Lite™ iFluor® 568 Tris NTA-Ni Complex (AAT Bioquest, Cat. No. 12617) with an additional 500 μL assay buffer (50 mM Tris-HCl pH 7.5, 0.1% Triton X-100 and 0.01% BSA). The reaction mixture was dispensed in a low volume 384-well white NBS plate (Corning, #3824) with 20 μL/well. Degrader molecules dissolved in DMSO were micro-dispensed with Multidrop™ Pico 8 Digital Dispenser (Thermo Fisher Scientific) to achieve serial dilution. After 2 h incubation at room temperature in the dark, 5 μM furimazine were dispensed with Pico 8 into each well and incubate for 3 min. Dual channel signal intensities were simultaneously measured on a BMG PheraStar plate reader mounted with 450±80nm and 610nm LP filter set, with auto gain setting on both channels, in precision mode.

### RNA extraction, Reverse Transcription qPCR

RNA for qPCR was extracted from cells using the RNeasy Plus Mini Kit (Qiagen, 74136) according to the manufacturer’s protocol. For quantification of VAV1 mRNA levels after treatment, expression of target genes was quantified using FlysisAmp Cells-to-CT 1-Step SYBR Green Kit (Vazyme, CL132-01). RT-qPCR was done on LightCycler® 480 Instrument II (384-multiwell plates). The following primers were used: VAV1 F TCAGTGCGTGAACGAGGTCAAG, VAV1 R CCATAGTGAGCCAGAGACTGGT, GAPDH F GGAGCGAGATCCCTCCAAAAT and GAPDH R GGCTGTTGTCATACTTCTCATGG (reference gene).

### Primary human T cells activation inhibition assay

Primary human T cells were obtained from peripheral blood mononuclear cells (PBMCs) sourced from the Gulf Coast Regional Blood Center. CD3+ T cells were isolated from PBMCs by negative selection using the magnetic bead–based MojoSort^TM^ Human CD3+ T Cell Isolation Kit (BioLegend, Cat. No. 480013). Briefly, 1× 10^8^ PBMCs were incubated with 100 μL of Biotin-Antibody Cocktail for 15 minutes on ice, followed by incubation with 100 μL of Streptavidin Nanobeads for 15 minutes. Unlabeled T cells were collected as the non-adherent fraction following magnetic separation. The purity of the isolated T cells was routinely confirmed by flow cytometry (BD, LSRII) using CD3 (BioLegend, Cat. No. 981012, Lot B438864) and CD19 (BioLegend, Cat. No. 982420, Lot B427354) lineage-specific markers. Isolated primary human T cells were plated in a 24-well plate at a density of 5 ×10^5^ cells/well. Cells were then treated with a 5-point, 5-fold serial dilution of the compound NGT-201-18 (starting concentration: 1000 nM) or DMSO vehicle control for 24 hours. Following compound treatment, the cells were activated using anti-CD3/CD28 magnetic beads for an additional 12 hours. Cellular analyses were subsequently performed by Western blotting for VAV1 and flow cytometry for the activation marker CD69. For flow cytometry, cells were harvested by centrifugation (500×g for 5 minutes) and resuspended in FACS buffer (PBS + 2% FBS). Cells were stained with PE anti-human CD69 Antibody (BioLegend, Cat. No. 310906, Lot B429659, 1:100 dilution) and DAPI (BioLegend, Cat. No. 422801, 30 μM) for dead cell exclusion, incubating on ice for 30 minutes. After one wash in FACS buffer, cells were centrifuged and subjected to analysis on the BD LSRII. Data was analyzed using FlowJo software.

### GluePlex workflow for VAV1:NGT-201-12:CRBN complex

To accurately predict the ternary complex structure of CRBN, VAV1, and the molecular glue, we developed and employed GluePlex, a multi-stage integrative modeling pipeline that combines geometric deep learning, physics-based docking, and templated co-folding.

Initial structural models for the VAV1 SH3-2 domain (UniProt: P15498, residues 782-842) and CRBN (UniProt: Q96SW2, residues 46-442) were retrieved from AlphaFold2 database. To identify potential PPI interfaces independent of the ligand, we utilized PeSTo (Protein Structure Transformer), a geometric deep learning model ^36^. PeSTo was applied to the unbound domain structures to predict interfacial residues with high PPI probability. Residues with a predicted interface probability score >0.3 were defined as “active residues” for downstream docking restraints. For CRBN, residues at DDB1 interfaces were excluded from the active residue list.

We then performed restrained, physics-based protein-protein docking using HADDOCK 2.4 (High Ambiguity Driven protein-protein DOCKing) ^37,46^ to sample the conformational space of the CRBN-VAV1 binary interaction. The active residues identified by PeSTo were used to define ambiguous interaction restraints (AIRs), forcing the docking algorithm to prioritize orientations where the predicted hotspots of VAV1 and CRBN engage. For both CRBN and VAV1, passive residues were auto-determined. All the other docking parameters are default. The resulting docked models were grouped into clusters based on interface root-mean-square deviation (i-RMSD) and ranked by HADDOCK score. One top-scored representative structure from each docking clusters (e.g., Clusters 1-6) were selected as templates for the subsequent co-folding stage.

To resolve the atomic details of the ligand-mediated interface, we utilized Boltz-2.2.1 ^38^ for templated co-folding. The docking result pdb files from each HADDOCK cluster were converted into mmCIF coordinates using ChimeraX 1.9. The mmCIF coordinates were supplied as a structural template for each Boltz-2 run, while the chemical structure of the molecular glue (e.g., NGT-201-12) was provided as a SMILES string. Each Boltz-2 run generates 20 different diffusion samples, and msa was disabled by explicitly stating “msa: empty” in the yaml files. For 6 HADDOCK clusters, a total of 120 co-folding structures were generated within 10 min on one NVIDIA H100 GPU.

The final outputs were visualized via UMAP ^47^ based on SH3-2 domain C-alpha atom coordinates after aligning to CRBN. Model selection was driven by the chainpair_ipTM (interface predicted TM-score) between VAV1 chain and CRBN chain, with the highest-scoring structure selected for further mechanistic analysis and mutagenesis validation.

### Molecular dynamics simulation

Molecular dynamics (MD) simulations were performed using the Desmond MD engine (Schrödinger, LLC) within Maestro 14.2. The initial ternary complex model was from GluePlex with NGT-201-12 positioned in the binding site. The initial binary complex model was generated by aligning (R)-enantiomers of NGT-201-17/18 and NGT-201-11/12 into the cereblon (CRBN) structure (PDB ID: 4CI3), excluding the DDB1 subunit. Protein-ligand complexes were prepared using the Protein Preparation Wizard in Maestro, which included removal of crystallographic water molecules, addition of hydrogen atoms, and restrained energy minimization.

The systems were solvated using the SPC water model in an orthorhombic box with a buffer distance sufficient to avoid periodic image interactions. Sodium and chloride ions were added to neutralize the system at a final ionic strength of 0.15 M. The OPLS4 force field was applied throughout all simulations.

A multi-stage equilibration protocol was performed prior to the production MD run. Stage 1 involved system building and setup. In Stage 2, Brownian dynamics simulations were conducted under the NVT ensemble at 10 K with reduced timesteps and positional restraints on solute heavy atoms (100 ps). This was followed by additional NVT equilibration at 10 K with restraints (12 ps, Stage 3), and two subsequent NPT equilibration stages also at 10 K with restraints maintained (12 ps each, Stages 4–5). In Stage 6, NPT equilibration was continued with all restraints removed (24 ps). Stage 7 constituted the 500 ns production run in the NPT ensemble at 300 K and 1.01325 bar, using the Martyna-Tobias-Klein barostat (τ = 2.0 ps) and Nosé-Hoover thermostat (τ = 1.0 ps). A multiple time-stepping scheme was applied with bonded, near, and far interactions evaluated every 0.002, 0.002, and 0.006 ps, respectively.

Simulation outputs included trajectory files recorded every 500 ps in DTR format (1000 frames per file), checkpoint files every 500 ps, and energy logs every 1.2 ps. Structure snapshots were saved every 500 ps with periodic boundary condition corrections. Initial velocities were assigned using a Maxwell-Boltzmann distribution at 300 K with a fixed seed for reproducibility. No restraints, enhanced sampling, or biasing potentials were applied during production. Final analyses, including ligand RMSD and torsional stability, were conducted using Desmond’s pl_analysis tools.

### Schrodinger Glide Docking and FEP+

For Glide Docking, GluePlex results were prepared with Schrodinger Protein Preparation Workflow and converted into docking grid. All selected ligands were prepared with Schrodinger LigPrep. Prepared ligands were docked into their corresponding docking grid with extra precision (XP), and the lowest docking scores were collected for comparison. Docked poses were rescored using GNINA v1.0 ^48^, yielding two scores per compound: CNNaffinity (CNN predicted binding affinity, pKd units) and Vinardo (empirical scoring function).

For FEP+, we selected NGT-201-18 GluePlex prediction structure as the receptor for simulation. With similar protocol previously mentioned, prepared ligands were docked into NGT-201-18 receptor grid with predicted NGT-201-18 as the reference ligand. The final docked structure with NGT-201-18 receptor and selected prepared ligands were used for FEP+ calculation. Three rounds of perturbations were carried out to calculate the final relative cooperativity with Equation 2 (A: CRBN, L: Molecular Glue, B: VAV1 SH3 domain) ^26^:

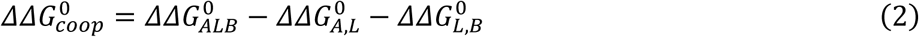

For binary energy predictions (ΔΔ*G*^0^*_A_*,*_L_*, ΔΔ*G*^0^*_L_*,*_B_*), we removed the other protein and use the structure for FEP+. When building the map for simulation, we selected NGT-201-11 and NGT-201-12 as biased nodes, for their representative structural features in our tested compounds. Default perturbation map from Schrodinger FEP+ was modified to increase connectivity of the map and accuracy of the perturbation. FEP+ calculation was submitted to Schrodinger Web Services.

For OpenFE calculations, we used the OpenFE (v1.8.0) framework. To ensure high-quality atom alignments, we employed the KartografAtomMapper with hydrogen mapping enabled for rigorous alignment across the chemical series. The transformation topology was defined using the generate minimal redundant network algorithm, which introduces strategic redundancy to facilitate cycle closure analysis. This design allows for the estimation of internal consistency and the determination of statistical uncertainties through cycle closure errors. Individual transitions were prioritized based on default LOMAP scores.

Alchemical simulations were performed using a Hamiltonian Replica Exchange approach at 298.15 K. Systems were solvated in a TIP3P water box and neutralized with 0.15 M NaCl. A 4-fs integration time step was utilized, enabled by hydrogen mass repartitioning to 3.0 amu and rigid water/HBonds constraints. Protein and solvent interactions were parameterized using a combination of Amber ff14SB, phosaa10, tip3p_standard, and tip3p_HFE_multivalent force fields, while ligands were modeled with openff-2.2.1. Electrostatics were treated via the Particle Mesh Ewald method with a 0.9 nm non-bonded cutoff.

The production protocol involved 5,000 steps of energy minimization followed by 1.0 ns of NPT equilibration. For each of the 11 replicas, 5.0 ns of production MD was conducted. Replica exchanges were attempted every 2.5 ps to ensure efficient sampling across the alchemical path. To handle vanishing and appearing atoms, we employed a Gapsys-type softcore LJ function (α = 0.85). Final free energy values and their associated standard errors were derived through a maximum likelihood estimator that incorporates the redundant cycle information from the network.

Absolute free energies of binding (AQFEP) ^49^ were computed using SandboxAQ’s internal platform. Each compound undergoes two independent free energy legs, bound (ligand in the VAV1 ternary complex) and unbound (ligand in solvent), with the absolute binding free energy computed as ΔG_binding = ΔG_bound − ΔG_unbound. Convergence is assessed independently for each leg using MBAR statistical overlap analysis. Uncertainty values are well within the range typically considered reliable for AFEP calculations, where errors below ∼1 kcal/mol are generally sufficient to correctly rank compounds.

### LIMD1 G-loop modeling

The G loop (residue 630-636) on the AlphaFold2 structure of LIMD1 LIM zinc binding domain 3 (residue 597-676) was aligned with CK1α G loop (residue 36-42) from PDB: 5FQD using the tmalign function in PyMOL. The protein complex was then loaded into Maestro 14.2 (Schrödinger, LLC) and prepared using protein preparation workflow to resolve clashes and perform local minimization. A 20 Å docking receptor grid was generated centered at the CRBN binding pocket. (R)-NGT-201-18 was prepared using LigPrep with OPLS4 forcefield and ligand docking was performed using the prepared molecule and receptor grid in XP precision by default settings.

### Dihedral angle potential energy surface scanning

Potential energy surface (PES) scans of key dihedral angles were performed using Gaussian 16 (Gaussian, Inc.) running on the Texas Advanced Computing Center (TACC) Frontera high-performance computing cluster. Initial geometries were first optimized using density functional theory (DFT) at the B3LYP level of theory with the 6-31+G(d,p) basis set. Solvent effects were modeled using the polarizable continuum model (PCM) for water, and Grimme’s D3 dispersion correction was applied to account for long-range van der Waals interactions. Following geometry optimization, relaxed dihedral scans were carried out using the Opt=ModRedundant keyword to systematically rotate the bond between the glutarimide ring and the adjacent phenyl ring (bond 1), plus the bond between two phenyl groups (bond 2) in 2D scanning. The dihedral angle was scanned in 10° increments over a full 360° rotation for bond 1, and 10° increments over a 180° rotation for bond 2, allowing the structure to relax at each fixed angle to identify the most energetically favorable conformations. Output log files were visualized using GaussView 5.0, and the resulting PES profiles were replotted in units of kcal/mol for downstream analysis.

### Reproducibility

For all the high-throughput proteomics experiments, at least two biological replicates with two MS technical replicates were analyzed for each treatment condition. All the biochemical experiments were completed with three biological replicates. Molecular dynamics simulation was performed in triplicate for 500 ns trajectory analysis.

## Data Availability

Chemical synthesis route and NMR spectra are provided as Supplementary Information. PDB structures used for modeling or scoring in this research can be found at 4CI3 [http://doi.org/10.2210/pdb4CI3/pdb], 5FQD [http://doi.org/10.2210/pdb5FQD/pdb], VAV1_AF2 [https://alphafold.ebi.ac.uk/entry/AF-P15498-F1], LIMD1_AF2 [https://alphafold.ebi.ac.uk/entry/AF-Q9UGP4-F1]. Modeled ternary complex of VAV1 and LIMD1 in PDB format are provided as supplementary data. The atomic coordinates of NGT-201-11/13/15/17 before and after DFT optimization and calculations are provided as supplementary data together with input settings and raw output energies and logs.

MS raw files and peak list mzML files are accessible through MassIVE database MSV000097850. [https://massive.ucsd.edu/ProteoSAFe/dataset.jsp?accession=MSV000097850] DIA-NN search results for global proteomics are available as supplementary data. All other raw data are available in the Source Data file.

## Code Availability

Hotspots on VAV1 and CRBN were calculated using web-based PeSTO (https://pesto.epfl.ch/). Restraint protein-protein docking was performed on the HADDOCK 2.4 server (https://rascar.science.uu.nl/haddock2.4/).

Codes for running Boltz-2 templated co-folding and UMAP analysis as part of the GluePlex protocol are available at https://github.com/Hanfeng-Lin/VAV1_GluePlex. The specific version of the code associated with this publication is archived in Zenodo and is accessible via 10.5281/zenodo.21306979 ^50^.

Codes for proteomics data analysis (associated with Supplementary Data 3) can be found at https://github.com/Hanfeng-Lin/VAV1_proteomics. The specific version of the code associated with this publication is archived in Zenodo and is accessible via 10.5281/zenodo.21306958 ^51^.

Code repositories generated in this research are all under MIT license.

## Acknowledgements

DFT calculations were performed on the Frontera system of Texas Advanced Computing Center (CHE25008). Desmond MD, Glide docking, and PyMOL visualization were carried out using the SBGrid software environment, which provides curated, version-controlled access to structural biology software and manages dependencies to support reproducible research. Software execution was supported through the SBGrid Capsules framework, which standardizes application environments and workflows across platforms, as described in ^52^.

## Author Contributions Statement

H.L. and J.W. conceptualized the study. H.L., X.Y., H.Z., R.C., M.Z., X.Q., Y.Y.Y., S.Z., R.Q., L.L., A.B. conducted experiments. All authors contributed to the development or design of methodology. H.L. and H.Z. developed the code, supporting algorithms and curated the data and repositories. H.L., X.Y., H.Z., M.Z., S.Z., R.Q., S.Y., analyzed the data. H.L. and J.W. drafted the manuscript. R.Q., A.N., X.C., J.W. guided and reviewed the manuscript. All authors read and approved of the final manuscript.

## References

1. Park, J., Cho, J. & Song, E. J. Ubiquitin-proteasome system (UPS) as a target for anticancer treatment. Arch. Pharm. Res. 43, 1144–1161 (2020).

2. Mullard, A. Targeted protein degraders crowd into the clinic. Nat. Rev. Drug Discov. 20, 247–250 (2021).

3. Fisher, S. L. & Phillips, A. J. Targeted protein degradation and the enzymology of degraders. Curr. Opin. Chem. Biol. 44, 47–55 (2018).

4. Burslem, G. M. et al. The Advantages of Targeted Protein Degradation Over Inhibition: An RTK Case Study. Cell Chem. Biol. 25, 67–77.e3 (2018).

5. Békés, M., Langley, D. R. & Crews, C. M. PROTAC targeted protein degraders: the past is prologue. Nat. Rev. Drug Discov. 21, 181–200 (2022).

6. Tsai, J. M., Nowak, R. P., Ebert, B. L. & Fischer, E. S. Targeted protein degradation: from mechanisms to clinic. Nat. Rev. Mol. Cell Biol. 25, 740–757 (2024).

7. Garber, K. The PROTAC gold rush. Nat. Biotechnol. 10.1038/s41587-021-01173-2 (2021) doi:10.1038/s41587-021-01173-2.

8. Oleinikovas, V., Gainza, P., Ryckmans, T., Fasching, B. & Thomä, N. H. From Thalidomide to Rational Molecular Glue Design for Targeted Protein Degradation. Annu. Rev. Pharmacol. Toxicol. 64, 291–312 (2024).

9. Krönke, J. et al. Lenalidomide causes selective degradation of IKZF1 and IKZF3 in multiple myeloma cells. Science 343, 301–305 (2014).

10. Petzold, G., Fischer, E. S. & Thomä, N. H. Structural basis of lenalidomide-induced CK1α degradation by the CRL4(CRBN) ubiquitin ligase. Nature 532, 127–130 (2016).

11. Watson, E. R. et al. Molecular glue CELMoD compounds are regulators of cereblon conformation. Science 378, 549–553 (2022).

12. Rankovic, Z. et al. Unbiased mapping of cereblon neosubstrate landscape by high-throughput proteomics. 2024.10.18.618633 Preprint at 10.1101/2024.10.18.618633 (2024).

13. Cao, S. et al. Defining molecular glues with a dual-nanobody cannabidiol sensor. Nat. Commun. 13, 815 (2022).

14. Hanzl, A., Inghelram, C., Schmitt, S. & Thomä, N. H. Primed for degradation: How weak protein interactions enable molecular glue degraders. Curr. Opin. Struct. Biol. 92, 103052 (2025).

15. Ito, T. et al. Identification of a primary target of thalidomide teratogenicity. Science 327, 1345–1350 (2010).

16. Lu, G. et al. The myeloma drug lenalidomide promotes the cereblon-dependent destruction of Ikaros proteins. Science 343, 305–309 (2014).

17. Sievers, Q. L. et al. Defining the human C2H2 zinc finger degrome targeted by thalidomide analogs through CRBN. Science 362, eaat0572 (2018).

18. Baek, K. et al. Unveiling the hidden interactome of CRBN molecular glues with chemoproteomics. BioRxiv Prepr. Serv. Biol. 2024.09.11.612438 (2024) doi:10.1101/2024.09.11.612438.

19. Petzold, G. et al. Mining the CRBN target space redefines rules for molecular glue–induced neosubstrate recognition. Science 389, eadt6736 (2025).

20. Dong, G., Ding, Y., He, S. & Sheng, C. Molecular Glues for Targeted Protein Degradation: From Serendipity to Rational Discovery. J. Med. Chem. 64, 10606–10620 (2021).

21. Koduri, V. et al. Targeting oncoproteins with a positive selection assay for protein degraders. Sci. Adv. 7, eabd6263 (2021).

22. Mayor-Ruiz, C. et al. Rational discovery of molecular glue degraders via scalable chemical profiling. Nat. Chem. Biol. 16, 1199–1207 (2020).

23. Yamanaka, S. et al. A proximity biotinylation-based approach to identify protein-E3 ligase interactions induced by PROTACs and molecular glues. Nat. Commun. 13, 183 (2022).

24. Costacurta, M. et al. Mapping the IMiD-dependent cereblon interactome using BioID-proximity labelling. FEBS J. 291, 4892–4912 (2024).

25. Che, X. & Team, T. S. M. I. YDS-GlueFold: Surpassing AlphaFold 3-Type Models for Molecular Glue-Induced Ternary Complex Prediction. 2024.12.23.630090 Preprint at 10.1101/2024.12.23.630090 (2025).

26. Dudas, B., Athanasiou, C., Mobarec, J. C. & Rosta, E. Quantifying Cooperativity through Binding Free Energies in Molecular Glue Degraders. J. Chem. Theory Comput. 10.1021/acs.jctc.5c00064 (2025) doi:10.1021/acs.jctc.5c00064.

27. Shalom, B., Salaymeh, Y., Risling, M. & Katzav, S. Unraveling the Oncogenic Potential of VAV1 in Human Cancer: Lessons from Mouse Models. Cells 12, 1276 (2023).

28. Ksionda, O. et al. Mechanism and function of Vav1 localisation in TCR signalling. J. Cell Sci. 125, 5302–5314 (2012).

29. Neurath, M. F. & Berg, L. J. VAV1 as a putative therapeutic target in autoimmune and chronic inflammatory diseases. Trends Immunol. 45, 580–596 (2024).

30. Razidlo, G. L. et al. Targeting Pancreatic Cancer Metastasis by Inhibition of Vav1, a Driver of Tumor Cell Invasion. Cancer Res. 75, 2907–2915 (2015).

31. Lin, H. et al. Lysineless HiBiT and NanoLuc Tagging Systems as Alternative Tools for Monitoring Targeted Protein Degradation. ACS Med. Chem. Lett. 15, 1367–1375 (2024).

32. Min, J., et al. Phenyl-Glutarimides: Alternative Cereblon Binders for the Design of PROTACs. Angew. Chem. Int. Ed. 60, 26663–26670 (2021).

33. Norris, S. et al. Design and Synthesis of Novel Cereblon Binders for Use in Targeted Protein Degradation. J. Med. Chem. 66, 16388–16409 (2023).

34. Wang, B., Cao, S. & Zheng, N. Emerging strategies for prospective discovery of molecular glue degraders. Curr. Opin. Struct. Biol. 86, 102811 (2024).

35. Kozicka, Z. & Thomä, N. H. Haven’t got a glue: Protein surface variation for the design of molecular glue degraders. Cell Chem. Biol. 28, 1032–1047 (2021).

36. Krapp, L. F., Abriata, L. A., Cortés Rodriguez, F. & Dal Peraro, M. PeSTo: parameter-free geometric deep learning for accurate prediction of protein binding interfaces. Nat. Commun. 14, 2175 (2023).

37. Honorato, R. V. et al. The HADDOCK2.4 web server for integrative modeling of biomolecular complexes. Nat. Protoc. 19, 3219–3241 (2024).

38. Passaro, S. et al. Boltz-2: Towards Accurate and Efficient Binding Affinity Prediction. 2025.06.14.659707 Preprint at 10.1101/2025.06.14.659707 (2025).

39. Leung, C. S., Leung, S. S. F., Tirado-Rives, J. & Jorgensen, W. L. Methyl Effects on Protein–Ligand Binding. J. Med. Chem. 55, 4489–4500 (2012).

40. Angell, R. et al. Biphenyl amide p38 kinase inhibitors 3: Improvement of cellular and in vivo activity. Bioorg. Med. Chem. Lett. 18, 4428–4432 (2008).

41. Reist, M., Carrupt, P.-A., Francotte, E. & Testa, B. Chiral Inversion and Hydrolysis of Thalidomide: Mechanisms and Catalysis by Bases and Serum Albumin, and Chiral Stability of Teratogenic Metabolites. Chem. Res. Toxicol. 11, 1521–1528 (1998).

42. Hughes, C. S. et al. Ultrasensitive proteome analysis using paramagnetic bead technology. Mol. Syst. Biol. 10, 757 (2014).

43. Hughes, C. S. et al. Single-pot, solid-phase-enhanced sample preparation for proteomics experiments. Nat. Protoc. 14, 68–85 (2019).

44. Demichev, V., Messner, C. B., Vernardis, S. I., Lilley, K. S. & Ralser, M. DIA-NN: neural networks and interference correction enable deep proteome coverage in high throughput. Nat. Methods 17, 41–44 (2020).

45. Kistner, F., Grossmann, J. L., Sinn, L. R. & Demichev, V. QuantUMS: uncertainty minimisation enables confident quantification in proteomics. 2023.06.20.545604 Preprint at 10.1101/2023.06.20.545604 (2023).

46. Honorato, R. V. et al. Structural Biology in the Clouds: The WeNMR-EOSC Ecosystem. Front. Mol. Biosci. 8, (2021).

47. McInnes, L., Healy, J., Saul, N. & Großberger, L. UMAP: Uniform Manifold Approximation and Projection. J. Open Source Softw. 3, 861 (2018).

48. McNutt, A. T. et al. GNINA 1.0: molecular docking with deep learning. J. Cheminformatics 13, 43 (2021).

49. Crivelli-Decker, J. E. et al. Machine Learning Guided AQFEP: A Fast and Efficient Absolute Free Energy Perturbation Solution for Virtual Screening. J. Chem. Theory Comput. acs.jctc.4c00399 (2024) doi:10.1021/acs.jctc.4c00399.

50. Hanfeng Lin. Hanfeng-Lin/VAV1_GluePlex: Publication_Ver. Zenodo 10.5281/ZENODO.21306979 (2026).

51. Hanfeng Lin. Hanfeng-Lin/VAV1_proteomics: Publication_Ver. Zenodo 10.5281/ZENODO.21306958 (2026).

52. Herre, C. et al. Introduction of the Capsules environment to support further growth of the SBGrid structural biology software collection. Acta Crystallogr. Sect. Struct. Biol. 80, 439–450 (2024).

